# RNA Polymerase III subunit Polr3a is required for craniofacial cartilage and bone development in zebrafish

**DOI:** 10.64898/2026.01.09.698267

**Authors:** Bailey T. Lubash, Roxana Gutierrez, Nicole A. Hansen, Kade Fink, Colette A. Hopkins, Lauren B. Sands, Jessica C. Nelson, Kristin E.N. Watt

**Affiliations:** Department of Craniofacial, Oral and Materials Sciences, School of Dental Medicine, University of Colorado Anschutz, Aurora, Colorado, United States of America; Department of Cell and Developmental Biology, School of Medicine, University of Colorado Anschutz, Aurora, Colorado, United States of America

## Abstract

Transcription by RNA Polymerase III (Pol III) is essential for ribosome biogenesis and translation in all cells, but pathogenic variants in genes encoding subunits of Pol III lead to tissue-specific phenotypes including craniofacial differences. To understand the function of Pol III in craniofacial development, we examined *polr3a* mutant zebrafish. These mutants display hypoplasia of the neural crest cell-derived craniofacial cartilage and bone but, surprisingly, no significant changes were observed in neural crest cell proliferation or survival during embryogenesis. At larval stages, increased cell death was observed throughout the head, including in the craniofacial cartilage. These changes coincide with reduced transcription of transfer RNAs and reduced ribosome biogenesis in *polr3a* mutant zebrafish. To determine tissue-specific transcriptional changes, we performed single-cell RNA-sequencing. Analysis revealed both global and cartilage-specific changes, including upregulation of *tp53*. However, Tp53 inhibition alone was not sufficient to rescue craniofacial cartilage and bone, indicating that additional factors are important to support cartilage and bone growth in *polr3a* mutants. Altogether, our study provides new mechanistic insights into the functions of Pol III in craniofacial development.

**Author Summary:** Craniofacial anomalies account for around 33% of all congenital birth defects and many of these are associated with defects in neural crest cell development. Disruptions in RNA Polymerase III, which plays important roles in ribosome biogenesis and translation, can result in tissue-specific phenotypes including craniofacial anomalies. However, the mechanisms underlying these craniofacial anomalies are not well understood. Here, we establish a zebrafish model to understand how a mutation in Pol III subunit *polr3a* affects craniofacial development. We observe hypoplasia of the craniofacial cartilage and bone in *polr3a* mutant zebrafish along with diminished transcription of transfer RNAs and reduced ribosome biogenesis. This leads to reduced proliferation and increased cell death throughout the head, but we do not detect any differences in early neural crest cell development. Using single-cell RNA-sequencing, we examine the transcriptional changes both broadly throughout the head and specifically within the craniofacial cartilage and identify upregulation of the Tp53 pathway. Inhibition of *tp53* only partially rescues cartilage and bone development, suggesting that Tp53-independent mechanisms contribute to cranioskeletal development in *polr3a* mutant zebrafish. Altogether, these studies highlight the critical function of Pol III during development and specifically in the differentiation and growth of craniofacial cartilage and bone.

## Introduction

Ribosome biogenesis is a global and essential cellular process that involves the coordinated synthesis of ribosomal RNA and ribosomal proteins. The 47S ribosomal RNA (rRNA) is transcribed by RNA Polymerase I (Pol I) in the nucleolus, which is then processed into the 18S, 28S, and 5.8S rRNAs while 5S rRNA is transcribed by RNA Polymerase III (Pol III). Pol III is a 17-subunit complex which transcribes several noncoding RNAs in addition to 5S rRNA, including transfer RNAs (tRNAs), 7SL RNA, and RMRP (encoding the RNA component of mitochondrial RNA processing endoribonuclease) [1]. 5S rRNA is a component of the 60S ribosomal subunit, while *RMRP* has functions in rRNA processing [2] and cell cycle progression [3], demonstrating that Pol III activity is important for ribosome biogenesis. In addition to this, Pol III transcribes *7SL,* the RNA component of the signal recognition particle which is responsible for co-translational targeting of secretory proteins to the endoplasmic reticulum [4], and tRNAs which are essential for translation. Thus, Pol III performs critical functions in the cell including roles in ribosome biogenesis and translation.

Structural studies have shown that POLR3A and POLR3B are the largest subunits of Pol III, forming the active site of the enzyme [5, 6]. Mice with mutations in Pol III subunit genes are embryonic lethal, demonstrating the essential function of Pol III during embryogenesis. Homozygous mutations in *Polr3b* are lethal prior to embryonic day (E) 9.5 [7], and null mutations in *Brf1*, which is a transcription factor required for Pol III-mediated transcription, are lethal prior to implantation (E3.5) [8]. Despite the essential function of Pol III, pathogenic variants in genes encoding subunits of Pol III are associated with several disorders, with many affecting neurological development [9, 10]. Pathogenic variants in *POLR3A* were first identified in individuals with POLR3-related leukodystrophy (POLR3-HLD; OMIM 607694) [11]. Clinical characteristics include hypomyelination in the central nervous system, motor dysfunction, abnormal dentition, and endocrine anomalies [12]. Craniofacial characteristics of individuals with POLR3-HLD include a flat midface, smooth philtrum, and pointed chin [13]. Further, several craniofacial differences including a small mandible were noted in a postnatal *Polr3b* conditional mutant mouse model, using a tamoxifen-inducible PDGFRα-Cre [14]. Distinct variants in *POLR3A* are associated with Wiedemann–Rautenstrauch syndrome (WRS; OMIM 264090), in which affected individuals display a triangular shaped face, pointed chin, and open cranial sutures [15, 16]. WRS and POLR3-HLD are inherited in an autosomal recessive manner, and both homozygous and compound heterozygous alleles have been described [11, 15–18]. Determining the genotype-phenotype relationship has been challenging for these POLR3-related disorders due to the large number of identified variants and combinations. Recent systematic analyses have demonstrated that variants associated with an individual disease do not show a high degree of spatial bias within the transcript or protein [16, 19].

Pathogenic variants in *POLR1C* and *POLR1D,* which form a heterodimer that is incorporated into the assembly platform for Pol I and Pol III, are associated with Treacher Collins syndrome (TCS; OMIM 248390 and 613717) [20]. Individuals with TCS show hypoplasia of the mandible, midface, as well as external and middle ears. Hypoplasia of the craniofacial cartilage and bone is recapitulated in *polr1c* and *polr1d* homozygous mutant zebrafish, demonstrating the importance of these subunits for normal craniofacial development [21]. Studies of a TCS variant in *POLR1D* in yeast demonstrated that it disrupts both Pol I and Pol III [22]. Additionally pathogenic variants in Pol III-associated transcription factors *BRF1* [23, 24] and *BRF2* [25] are all linked to craniofacial differences, which further suggests an important function for Pol III in craniofacial development. Disruptions in ribosome biogenesis lead to a group of syndromes known as ribosomopathies, of which Treacher Collins syndrome is one example.

Ribosomopathies display distinct and tissue-specific phenotypes, many of which affect craniofacial development [26, 27]. However, it should be noted that because of the variety of transcriptional changes that can occur as a result of disruption of Pol III, not all Pol III-related disorders are considered ribosomopathies. Altogether, this suggests that there is a critical function for Pol III in craniofacial development, as well as in other tissues, but when and how these craniofacial differences arise remain unknown.

Craniofacial development begins early in embryogenesis and requires coordination of the growth and development of multiple tissues. Neural crest cells (NCCs) are particularly important for craniofacial development, forming much of the cartilage, bone, and connective tissue in the head and face [28, 29]. The frequency of craniofacial differences in ribosomopathies has led to the hypothesis that NCCs are especially sensitive to disruptions in ribosome biogenesis, with ribosome biogenesis having an important role in NCC development and epithelial to mesenchymal transitions [30, 31]. Prior studies demonstrated that NCCs exhibit elevated rRNA transcription and translation relative to surrounding tissues during embryogenesis [30]. Further, cells with high levels of rRNA transcription show high levels of p53 stabilization and are more likely to trigger a cell death response [32]. Together, this suggests that NCCs are especially sensitive to perturbations in ribosome biogenesis and translation and more likely to undergo cell death. Consistent with this idea, *polr1c* and *polr1d* mutant zebrafish show increased cell death in NCC progenitors within the neuroepithelium [21]. Conditional deletion of Pol III subunits has provided information on the response of specific tissues to the loss of Pol III, including the intestine and brain [14, 33, 34], but our understanding of the effects of diminished Pol III during embryogenesis and NCC development remains incomplete.

Here, we assess the function of Pol III during NCC and craniofacial development by disrupting Polr3a, the largest Pol III subunit that forms the catalytic core of the enzyme. Polr3a is highly conserved in vertebrates, with 90% identity at the protein sequence level between zebrafish and humans. We established a *polr3a* mutant zebrafish and examined the development of NCC-derived craniofacial cartilage and bone. Homozygous mutant zebrafish are viable through 8 days post fertilization (dpf), circumventing the need for conditional deletion and enabling the study of early embryonic, global, and tissue-specific roles for *polr3a.* Surprisingly, we found that ubiquitous reduction in *polr3a* does not affect early NCC development, but that there is a significant effect on the growth and differentiation of craniofacial cartilage and bone after 3 dpf. This coincides with global reductions in tRNA and rRNA along with increased cell death and reduced proliferation. We performed single-cell RNA-sequencing and assessed the development of multiple tissues in *polr3a* mutant zebrafish. Some transcriptional changes, including *tp53* upregulation, are associated with only a subset of cells in the head, including the craniofacial cartilage. Altogether, these results demonstrate that zygotic Pol III is required after the initiation of craniofacial cartilage and bone development. The *polr3a* loss-of-function mutants will serve as useful models for furthering our understanding of *polr3a* during development and in the pathogenesis of POLR3-related disorders.

## Results

### Establishing a *polr3a* mutant zebrafish for the study of craniofacial development

Craniofacial phenotypes associated with disruption of Pol III in humans [13, 16] and mice [14] suggest that Pol III is required for the growth and development of the craniofacial skeleton. However, whether Pol III is required at earlier developmental stages for NCC development and differentiation into craniofacial cartilage and bone remained undetermined. To study the consequences of Pol III perturbation during craniofacial development, we examine zebrafish with a mutation in *polr3a*, encoding the largest subunit of Pol III (Fig. 1D). We obtained zebrafish with an ENU-induced nonsense mutation in exon 14 of *polr3a*, *polr3a^sa907^*, which results in a premature stop codon at amino acid 623 of 1390, located in the pore domain of the protein (Fig 1 A-C; S1 Fig). The sa907 allele occurs at a conserved residue near other variants identified in POLR3-related leukodystrophy and WRS (Fig 1 C; S1 Fig) and may serve as a useful disease model.

**Fig. 1.**
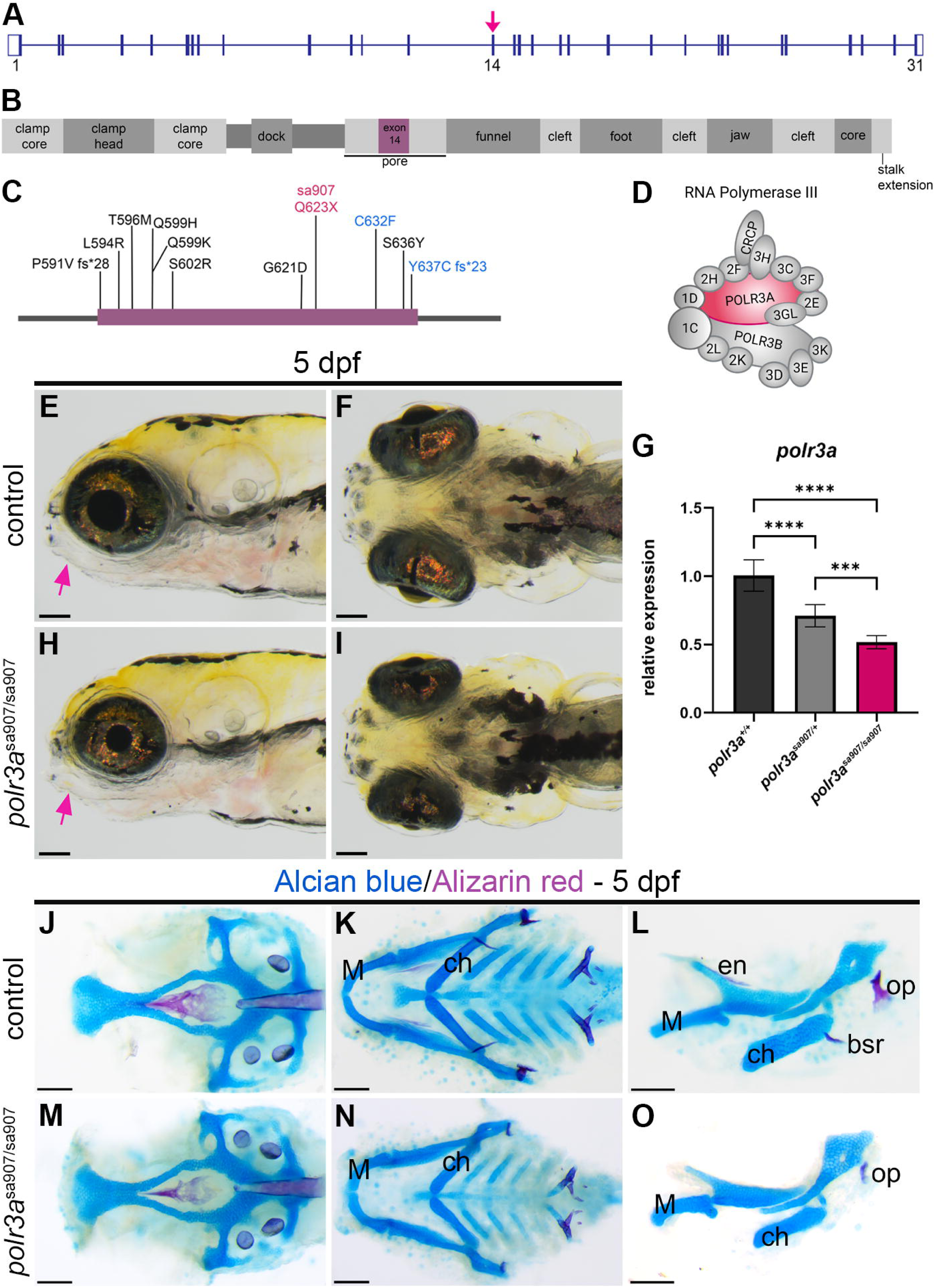
***polr3a^sa907^*disrupts craniofacial development.** (A) Representation of the exons of *polr3a* in zebrafish. The sa907 allele is in exon 14 (magenta arrow). (B) Schematic of protein domains of human POLR3A (based on Girbig *et al.,* 2021). The amino acid sequence encoded by exon 14 is highlighted in purple, located in the pore domain. (C) Location of pathogenic variants within the protein domain encoded by exon 14 associated with POLR3-related leukodystrophy (black), WRS (blue), and the location of sa907 (magenta). (D) Representation of the 17-subunit RNA Polymerase III complex highlighting POLR3A, magenta. (E-I) Brightfield images at 5 dpf shows an overall change in the size of the head, including the jaw (arrows; E,H). Ventral views (F, I) show the overall length and width of the head is reduced. (G) qRT-PCR shows that the sa907 allele results in a significant reduction in *polr3a* mRNA expression in *polr3a^sa907/sa907^*homozygous mutant zebrafish at 3 dpf, and a slight reduction in *polr3a^sa907/+^*heterozygous zebrafish. ****p < 0.0001; ***p = 0.0005. (J-O) Alcian blue and Alizarin red staining shows overall reduction in the elements of the viscerocranium (K, N). Reductions in cartilage and bone elements including the opercle (op), Meckel’s cartilage (M), and the ceratohyal (ch), entopterygoid (en), and branchiostegal rays (bsr) are apparent by 5 dpf (H, K). Scale bar = 100 µm. Panel D Created in BioRender. Watt, K. (2026) https://BioRender.com/j5w61o1.

We examined *polr3a^sa907/sa907^* zebrafish at 5 dpf and observed overall changes in head development, including the developing jaw (Fig 1 E-I, arrows). Quantitative RT-PCR (qRT-PCR) for *polr3a* in *polr3a^sa907/sa907^*mutants and control siblings showed that there is a significant decrease in *polr3a* in homozygous mutants as well as a slight decrease in heterozygous animals at 3 dpf (Fig 1 G). Given the ∼50% decrease in transcript levels, we predict that sa907 leads to some nonsense mediated decay. At the protein level, we detected very little full length Polr3a protein (S1 Fig). It is possible a small amount of truncated protein is produced, as we also detect a band near 78 kDa, which is the predicted size of the mutant protein (S1 Fig). Next, to determine the effects of reduced Polr3a on cranioskeletal development, we performed an Alcian blue and Alizarin red stain for cartilage and bone, respectively (Fig 1 J-O). While all craniofacial cartilage elements were present, they were reduced in size in *polr3a^sa907/sa907^* zebrafish compared to control siblings (Fig 1 L vs. O; S2 Fig). The opercle, branchiostegal rays, and entopterygoid bones were either reduced in size or absent in *polr3a^sa907/sa907^*zebrafish (Fig 1 L, O) These bones did not recover by 8 dpf (S3 Fig), suggesting that these differences are not simply due to developmental delay. The *polr3a^sa907/sa907^* zebrafish are not viable after 8 dpf, preventing examination of later stages of bone development. These data demonstrate that Polr3a is required for the development and growth of the NCC-derived craniofacial skeleton in zebrafish.

### Craniofacial cartilage and bone elements are hypoplastic in *polr3a^sa907/sa907^* zebrafish

We hypothesized that Pol III would be critical during NCC development as NCCs contribute to the majority of the craniofacial cartilage and bone, and NCCs undergo significant shifts in rRNA transcription during their development [30, 31]. To determine the onset of differences in *polr3a* mutant zebrafish, we examined zebrafish from 36 hours post fertilization (hpf) through 5 dpf. We assessed the NCC population and cells within the NCC lineage using the transgenic reporter *Tg(-7.2sox10:egfp)* [35], referred to as *sox10:egfp*. At 36 hpf, when NCCs have migrated into the pharyngeal arches, we did not observe any significant differences between control and *polr3a^sa907/sa907^* mutants and the pharyngeal arches are the same size (S4 Fig A-C). This suggests that early NCC development proceeds normally. Examination of *sox10:egfp* expression and quantification of the pharyngeal arches at 48 hpf also showed no significant differences (S4 Fig D-F) and no differences were detected in the overall morphology of the embryos.

Consistent with this, the pharyngeal arch domain appeared similar using in situ hybridization for *dlx2a* (S5 Fig), which is expressed in the NCCs in the pharyngeal arches [36]. Assessment of *sox10:egfp* expression at 3, 4, and 5 dpf did not reveal significant alteration in the domain of expression (Fig 2 A-F), although both the overall head and cartilage elements appeared smaller in size by 4 dpf.

**Fig 2.**
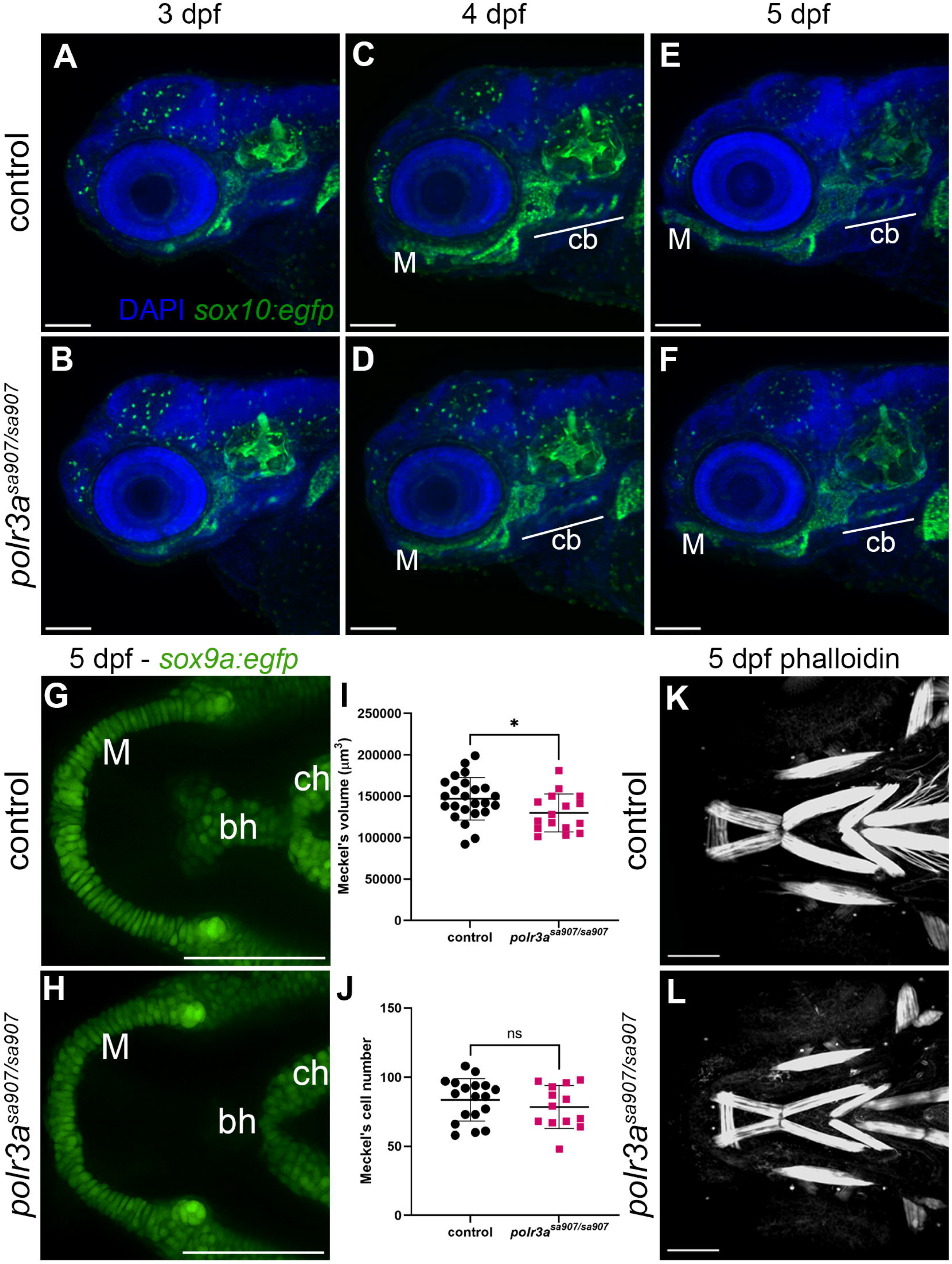
Neural crest cell-derived craniofacial cartilage is reduced in *polr3a^sa907/sa907^* zebrafish. (A-F) Examination of the craniofacial cartilage using *sox10:egfp* (green) and all nuclei (DAPI, blue) shows a reduction in overall head size and cartilage elements at 5 dpf and 4 dpf, including Meckel’s cartilage (M) and the ceratobranchials (cb). No changes were detected at 3 dpf and mutants appear phenotypically normal. 3 dpf: n = 12 controls, 12 mutants; 4 dpf: n = 12 controls 12 mutants; 5 dpf: n = 17 controls, 16 mutants. (G,H) Cartilage was examined at 5 dpf using *sox9a:egfp*. A reduction in size is evident for in the basihyal (bh), while Meckel’s cartilage (M) shows a change in morphology. ch, ceratohyal. (I) Quantification shows that the overall size of Meckel’s is reduced (*p = 0.035). (J) Cell number within Meckel’s cartilage was unchanged (ns, p = 0.367). n = 18 controls, 13 mutants. (K, L) Phalloidin staining at 5 dpf shows that muscle patterning is normal, however, the overall size of the muscles appears to be reduced in size consistent with an overall reduction in head size. Scale bar = 100 µm.

To better visualize changes in cartilage morphology, we used the transgenic reporter *sox9a:egfp* [37]. Cartilage elements such as Meckel’s cartilage appear shorter and wider by 5 dpf in *polr3a^sa907/sa907^* zebrafish and the basihyal was also reduced in size (Fig 2 G,H). Analysis of Meckel’s cartilage volume at 5 dpf revealed it was slightly smaller in mutants (Fig 2I). Despite the overall reduction in size of Meckel’s cartilage, we did not detect significant changes in cell number (Fig 2J). We next examined whether the average cell size was smaller or if there were changes in cell morphology that could account for this size difference. We found that the average area of the cells in Meckel’s cartilage were not significantly different between mutants and controls, nor was the eccentricity, which is a measure of how round versus columnar the cells appear (S6 Fig). However, this very slight change could be due to regional differences in the cells within Meckel’s cartilage, as we observed the cells near the midline appeared more disorganized in mutants relative to controls in our analysis of the *sox9a:egfp* as well as Alcian blue stains (Figs 1, S2, S3).

This disorganization led us to hypothesize that some of the differences in cartilage development and the morphology of Meckel’s cartilage in *polr3a^sa907/sa907^* zebrafish could be a consequence of disrupted muscle development, as muscle contractions are required for normal development of the cranioskeletal elements [38, 39]. Labeling muscles with Phalloidin staining at 6 dpf revealed that there were no significant changes in muscle patterning in the region of the developing jaw; however, the overall size appeared reduced, consistent with the overall smaller head of *polr3a^sa907/sa907^* mutants (Fig 2 K,L). The overall reduction in craniofacial muscle suggests that non-NCC-derived tissues are also affected in *polr3a^sa907/sa907^* mutants, consistent with a broad function for Pol III during development. Altogether, these data suggest that the phenotypic onset is after NCC migration and instead occurs during the differentiation and growth of the craniofacial skeleton.

As changes in morphology in *polr3a^sa907/sa907^* zebrafish were apparent by 4 dpf, we hypothesized that differences in cartilage and bone were a result of a change in differentiation. In situ hybridization for *col2a1a,* which is expressed in the developing craniofacial cartilage [40], showed no changes in the domain of expression within the developing cartilage at either 48 hpf or 72 hpf in *polr3a^sa907/sa907^* zebrafish (S7 Fig). We assessed whether there were any changes in bone development in *polr3a^sa907/sa907^* zebrafish using the transgenic reporter *Tg(sp7:gfp)* [41], which is expressed in the osteoblast population, in combination with Alizarin red staining for mineralized bone. We initially examined the domain of *sp7:gfp* expression and Alizarin red staining in the opercle, as the opercle is one of the first craniofacial bones to develop and its development has been well characterized in zebrafish [42]. Quantification of *sp7:gfp* in the opercle at 3 dpf did not show any significant differences between controls and *polr3a^sa907/sa907^*zebrafish (S8 Fig). This suggests that osteoblast specification is not affected at 3 dpf.

By 4 dpf, we detected significant reductions in the overall domain of *sp7:gfp* expression as well as Alizarin red in *polr3a^sa907/sa907^* zebrafish (Fig 3 A,B) and the opercle remains reduced in size through 6 dpf (Fig 3 C,D) and 8 dpf (S9 Fig). Differences in Alizarin red were not limited to the opercle, with reductions observed in nearly all craniofacial bones (Fig 3 I-N), Interestingly, the overall volume of *sp7:gfp* in the dentary was not significantly different in mutants (Fig 3 K), although the domain of *sp7:gfp* expression appears shorter and wider relative to controls (Fig 3 J). This suggests that the organization, but not overall osteoblast specification, may be affected in *polr3a* mutants. In contrast, Alizarin red within the dentary was significantly reduced (Fig 3 L). The maxilla showed reduced *sp7:gfp* and Alizarin red volume (Fig 3 M,N), as did the branchiostegal rays, demonstrating that all bones examined were reduced in size.

**Fig 3.**
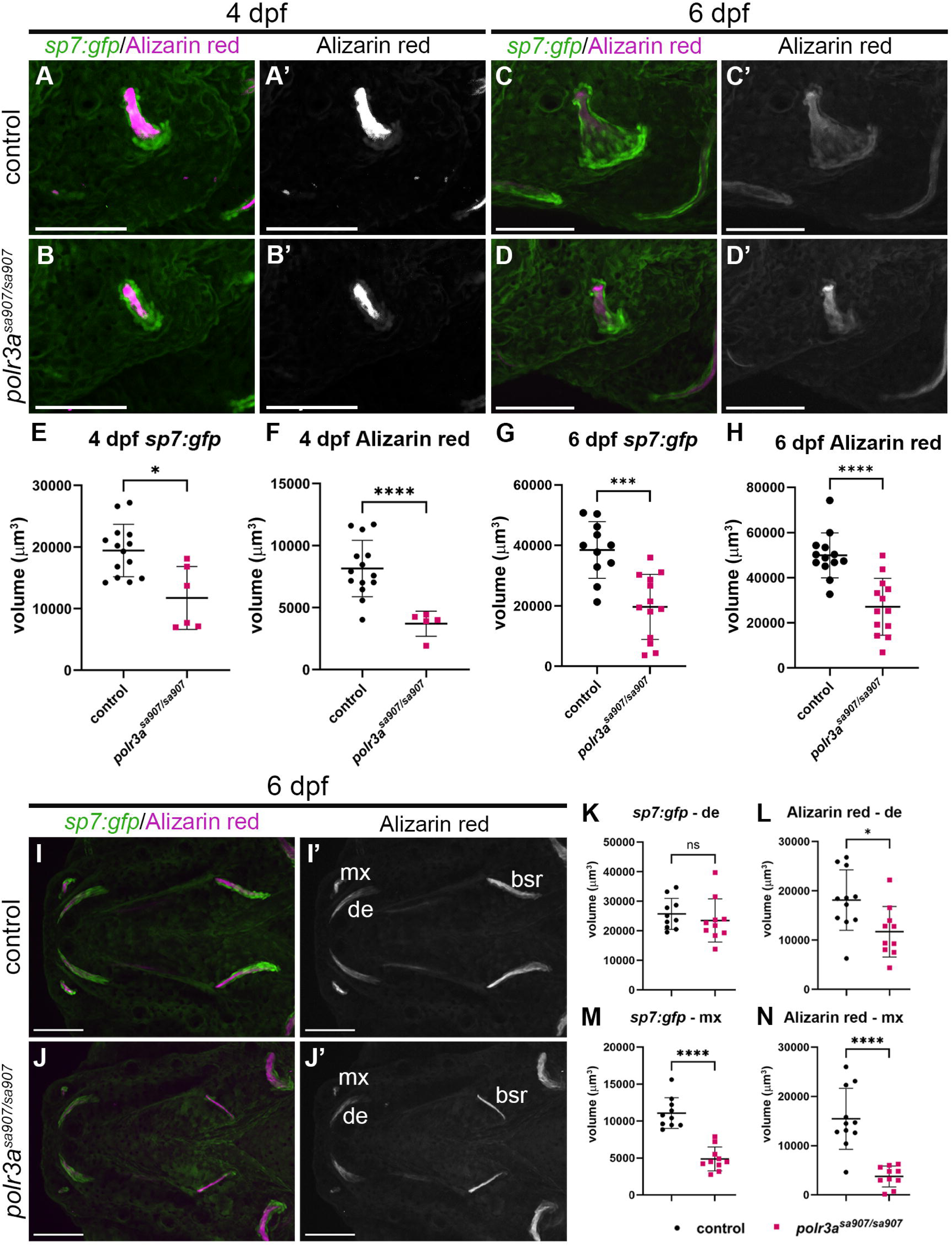
Bone elements are reduced in *polr3a^sa907/sa907^* zebrafish. (A,B) Images of the opercle at 4 dpf expressing *sp7:gfp* (osteoblasts) and stained with Alizarin red (bone, magenta) show that both the osteoblasts (E; *p = 0.01) and bone (F; ****p < 0.0001) are significantly reduced in mutants. n = 14 controls, 6 *polr3a^sa907/sa907^*. (C,D) Images of the opercle at 6 dpf show that it is significantly reduced in size and that osteoblasts (G; ***p = 0.0003) and bone (H; ****p < 0.0001) remain significantly reduced in mutants. n = 11 controls, 13 *polr3a^sa907/sa907^*. (I,J) Ventral views of control and *polr3a^sa907/sa907^* zebrafish reveals hypoplasia of multiple cranial bones including the maxilla (mx), dentary (de) and branchiostegal rays (bsr). (K,L) Quantification of *sp7:gfp* and Alizarin red within the dentary. The dentary is shorter and wider in *polr3a^sa907/sa907^* zebrafish, but no significant changes were observed in the overall volume (K; ns, p = 0.28) while Alizarin red is significantly reduced (L; *p = 0.01). The maxilla is smaller in size overall with reductions in both *sp7:gfp* (M; ****p < 0.0001) and Alizarin red (N; ****p < 0.0001). n = 11 controls, n = 10 *polr3a^sa907/sa907^*. Scale bar = 100 µm.

To determine if these differences persist, we quantified Alizarin red staining at 8 dpf. At the end of the lifespan of *polr3a* mutant larvae, all bones remain significantly reduced in size including the dentary, maxilla, and branchiostegal rays (S9 Fig). At 8 dpf, there is only a single branchiostegal ray and the opercle fails to expand to form a fan-like structure typically observed by 5 dpf. While we see some endochondral ossification in *polr3a^sa907/sa907^* zebrafish in the hyosymplectic, very little is detected in the ceratohyal (S9 Fig), indicating that both intramembranous and endochondral ossification are affected in *polr3a* mutant zebrafish. Altogether, these data led us to hypothesize that mineralization was affected in *polr3a^sa907/sa907^* zebrafish.

A potential contributing factor to reduced bone in cranioskeletal development is calcium homeostasis. A recent report showed that NCC-derived *sox10+* cells surround the corpuscles of Stannius (CS) in zebrafish, an endocrine gland which forms at the junction between the distal early tubule and the distal late segment of the pronephric tubules and releases the calcium regulatory hormone Stanniocalcin 1l [43]. Stanniocalcin 1l (*stc1l*) in turn suppresses calcium uptake, therefore playing a key role in regulating bone mineralization [44]. In *sox10* mutants, the CS is enlarged and more strongly expresses the anti-hypercalcemic hormone, *stc1l,* leading to a delay in mineralization [43]. To assess whether altered CS development and disrupted Ca^2+^ homeostasis might contribute to the skeletal hypoplasia in our mutants, we examined expression of *cdh16* at 5 dpf, which is expressed in the corpuscles from 2 dpf [45, 46]. We did not identify any significant differences in *cdh16* expression in homozygous mutants (S10 Fig A-C). Reduced calcium homeostasis also results in hypersensitivity to the acoustic startle response [46, 47], so we next assessed the sensitivity of *polr3a^sa907/sa907^* zebrafish. These studies showed that the larvae were hyposensitive to acoustic stimuli across two experimental replicates (S10 Fig D). *polr3a^sa907/sa907^*zebrafish also trended toward improved habituation to repeated acoustic stimuli (S10 Fig E). This effect was statistically significant in one experimental replicate for stimuli 21-30, but did not repeat. Habituation to the final stimuli (31-40) was unchanged across experimental replicates (S10 Fig F). This further suggests that differences in bone development are not due to a reduction in overall whole-body calcium metabolism, which we expect would cause hypersensitivity and reduced habituation. Altogether, these data demonstrate that there are differences in craniofacial cartilage and bone development in *polr3a^sa907/sa907^*mutants relative to controls, but we find no evidence that these differences are due to changes in the NCC population prior to 3 dpf or a result of NCC-dependent control of Ca^2+^-regulatory endocrine tissues.

### *polr3a* is broadly expressed during zebrafish embryogenesis

Pol III is globally required for ribosome biogenesis and translation, and therefore multiple tissues and structures are predicted to be disrupted in *polr3a^sa907/sa907^*mutants. Interestingly, *polr3a^sa907/sa907^* zebrafish are morphologically indistinguishable from controls prior to 4 dpf (Fig 2), which led us to hypothesize that maternal contribution of Pol III and *polr3a* may be sufficient for early development.

Maternal ribosomes and rRNA have been detected in zebrafish up to 4 dpf [48, 49] and are sufficient to support early embryonic development in Pol I mutant zebrafish [50, 51]. We performed qRT-PCR for *polr3a* from the 4-cell stage though 5 dpf in wild-type zebrafish (S11 Fig A). At the 4-cell stage, *polr3a* is expressed at a higher level relative to any other developmental stage, and diminishes in expression until 2 dpf, and increases again after 3 dpf. As a complementary approach, we performed *in situ* hybridization chain reaction (HCR) for *polr3a* in wild-type zebrafish at 24 hpf, 48 hpf, and 72 hpf (S11 Fig B-D). These data showed that *polr3a* is broadly expressed throughout the embryo, consistent with the expectation that Pol III is required globally. The relative expression of *polr3a* started increasing at 4 and 5 dpf, which is when *polr3a^sa907/sa907^*zebrafish begin to present phenotypic differences from controls. This suggests that there is considerable maternal expression of *polr3a* which may facilitate embryonic development at early stages, but this maternal expression is not sufficient to support the continued growth of the animals after 3 dpf.

We next hypothesized that observed differences in bone development (Fig 3) may occur because of the timing of the initiation of bone development around 3 dpf, which coincides with increased zygotic *polr3a* expression in zebrafish. However, this mechanism may not be unique to bone, as other tissues, such as myelin, also initiate development at these later stages [52]. Using expression of *Tg(mbp:tagRFP)* [53] as a marker of myelinating cells, we did not detect any significant changes in the overall domain of expression of *mbp* in *polr3a^sa907/sa907^*mutants in the head or trunk (S12 Fig). This is in contrast to the clear differences we observed in the osteoblast population (Fig 3) and raises the intriguing possibility that *polr3a* and tRNAs may be differentially required across tissues. After depletion of maternal stores, some tissues show significant changes, such as the craniofacial bone, while others, such as *mbp-*expressing cells, do not show differences based on our current analyses.

### *polr3a^sa907/sa907^* zebrafish display changes in proliferation and cell death

We next hypothesized that increased cell death and reduced proliferation contribute to changes in cartilage and bone development. We used TUNEL and an anti-phospho-histone H3 Ser 10 (pH3) antibody to quantify changes in cell death and proliferation, respectively. We examined embryos from 2 dpf, prior to any detectable phenotype, through 5 dpf, when mutant zebrafish are clearly identifiable based on an overall reduction in head size. To determine whether changes in cell death and proliferation affected craniofacial morphology specifically within the developing craniofacial cartilage, we utilized *sox10:egfp* to delineate the pharyngeal arch 1 and 2-derived cartilage and quantified pH3 and TUNEL specifically within this region. Consistent with our previous observations, there were no significant differences at 2 dpf (S13 Fig). However, by 3 dpf, there was a significant increase in cell death (Fig 4 A’ vs. B’, C), which is prior to any detectable differences in phenotype. By 4 dpf, we began to observe changes in the overall shape of cartilage elements including the basihyal and ceratohyal (Fig 4 E vs. F). Interestingly, cell death was unchanged at this stage which we predict is due to stress responses and an attempt to repair damage (see also Figs 6, 7). Cell death is again increased at 5 dpf (Fig 4 I’ vs. J’; K) with a notable increase within Meckel’s cartilage. Interestingly, no significant changes in pH3 were detected within the cartilage of *polr3a^sa907/sa907^* zebrafish at any stage (Fig 4; S13 Fig). It remains possible that there are changes in proliferation that were not detected using pH3, which labels cells in G2 to M phase [54]. However, as proliferation is not a major contributor to cartilage growth in zebrafish until after 4 dpf [55], this data suggests that differences in the size of cartilage elements may be due to increased cell death as opposed to differences in proliferation.

**Fig 4.**
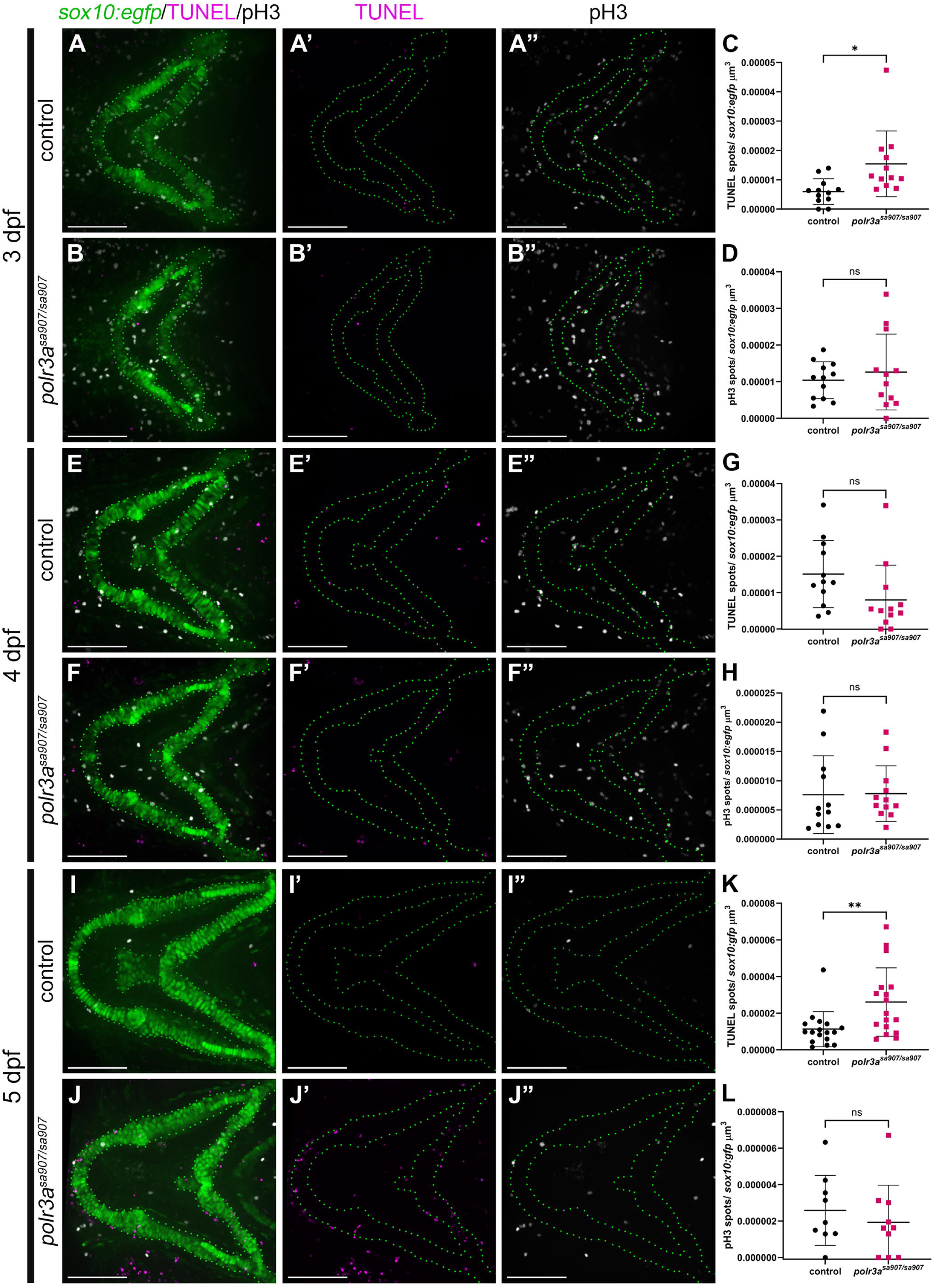
Cell death is increased within the craniofacial cartilage in *polr3a^sa907/sa907^* zebrafish. Cell death (TUNEL, magenta) and proliferation (pH3, white) were quantified within pharyngeal arch 1 and 2 derived cartilage elements using *sox10:egfp* (green, outlined) to mark this domain. (A,B) At 3 dpf, there is a significant increase in cell death (C; *p = 0.016), but no change in proliferation (D; ns, p = 0.51). n = 12 controls, 12 *polr3a^sa907/sa907^*. (E,F) At 4 dpf, differences in the morphology of the craniofacial cartilage are visible, including a reduced basihyal, while cell death (G; ns, p = 0.08) and pH3 remain unchanged (H; ns, p = 0.94). n = 12 controls, 12 *polr3a^sa907/sa907^*. (I,J) By 5 dpf, there is a noted difference in cartilage morphology and increased cell death (K; **p = 0.008; n = 17 controls, 17 *polr3a^sa907/sa907^*), while pH3 is unchanged (L; ns, p = 0.48; n = 9 controls, 10 *polr3a^sa907/sa907^*), suggesting that differences are due to changes in cell death. Scale bar = 100 µm.

To understand if increased cell death was specific to the craniofacial cartilage or occurred more broadly, we examined cell death and proliferation throughout the whole head. No significant changes were observed at 2 dpf (S14 Fig); however, at 3 dpf increased TUNEL staining was observed in *polr3a^sa907/sa907^* zebrafish, with areas near the eye and tectum consistently showing increased cell death (Fig 5 A’ vs. B’, C). At 4 and 5 dpf, cell death remains increased in *polr3a^sa907/sa907^* zebrafish (Fig 5 E’ vs. F’, quantification in G; I vs J, quantification in K). Quantification of pH3 revealed that pH3+ cells were significantly diminished in *polr3a^sa907/sa907^* zebrafish at 3 dpf and this trend persisted through 5 dpf (Fig 5 D, H, L). Altogether, this demonstrates that both increased cell death and reduced proliferation could contribute to the reduced head size of *polr3a^sa907/sa907^* zebrafish, while increased cell death may be driving differences in the craniofacial cartilage within *polr3a^sa907/sa907^* zebrafish.

**Fig 5.**
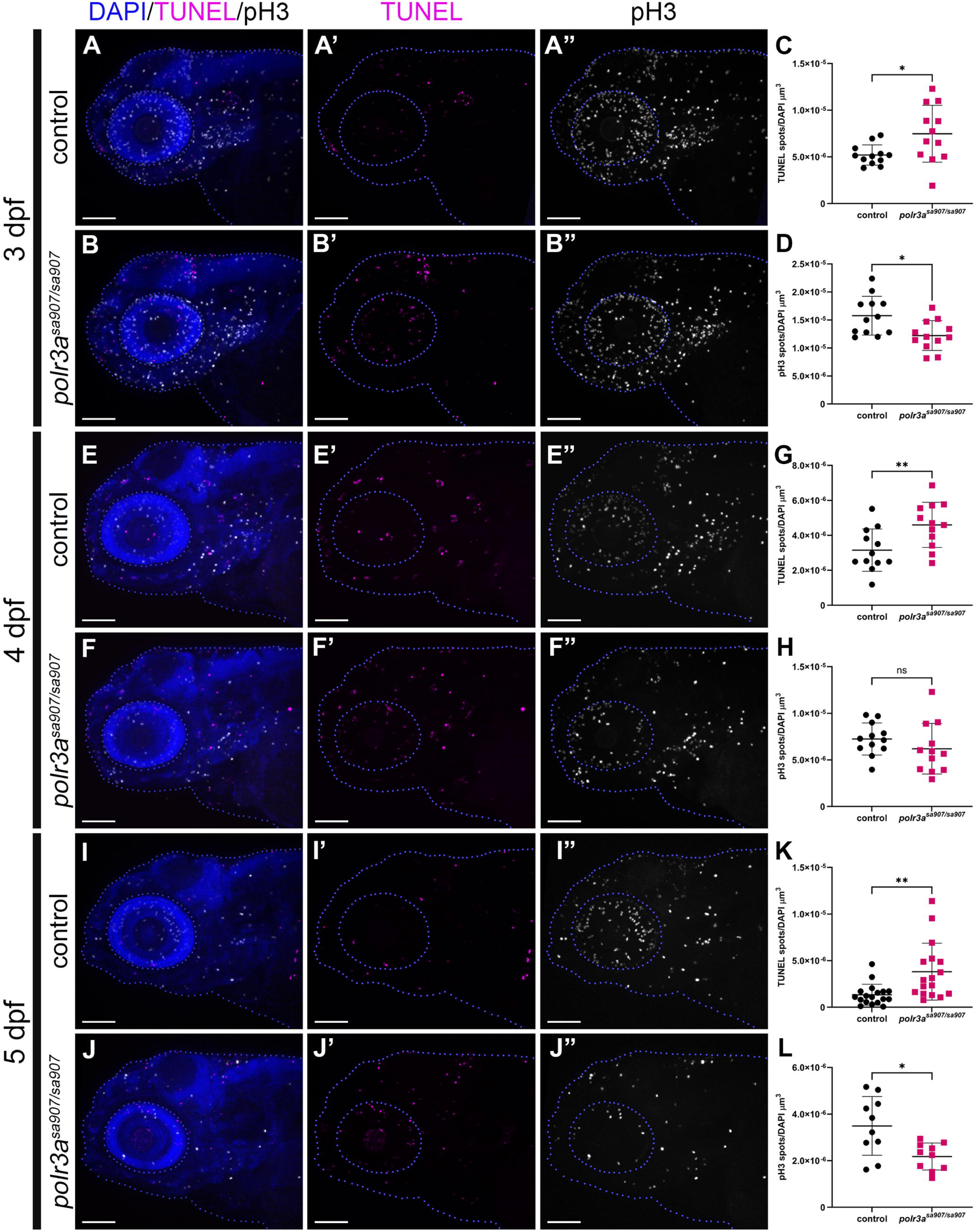
*polr3a^sa907/sa907^* zebrafish exhibit reduced globally proliferation and increased cell death. TUNEL staining for cell death (magenta) and pH3 staining for proliferation (white) from 3 – 5 dpf were quantified throughout the cranial region and normalized to DAPI (blue, outlined). (A, B) At 3 dpf, increased cell death was observed in the tectum and eye along with a slight decrease in proliferation. TUNEL is significantly increased (C; *p = 0.03) while pH3 is significantly decreased (D; *p = 0.01). n = 12 controls, n=12 *polr3a^sa907/sa907^*(E,F) At 4 dpf, cell death remains increased (G; **p = 0.0098), and pH3 was slightly reduced, but this was not statistically significant (H; ns, p = 0.27). n = 12 controls, n = 12 *polr3a^sa907/sa907^*. (I,J) By 5 dpf TUNEL remains significantly increased (K; **p = 0.005) and pH3 is significantly reduced (L; *p = 0.01). n = 17 controls, n = 17 *polr3a^sa907/sa907^*. Scale bar = 100 µm.

**Fig 6.**
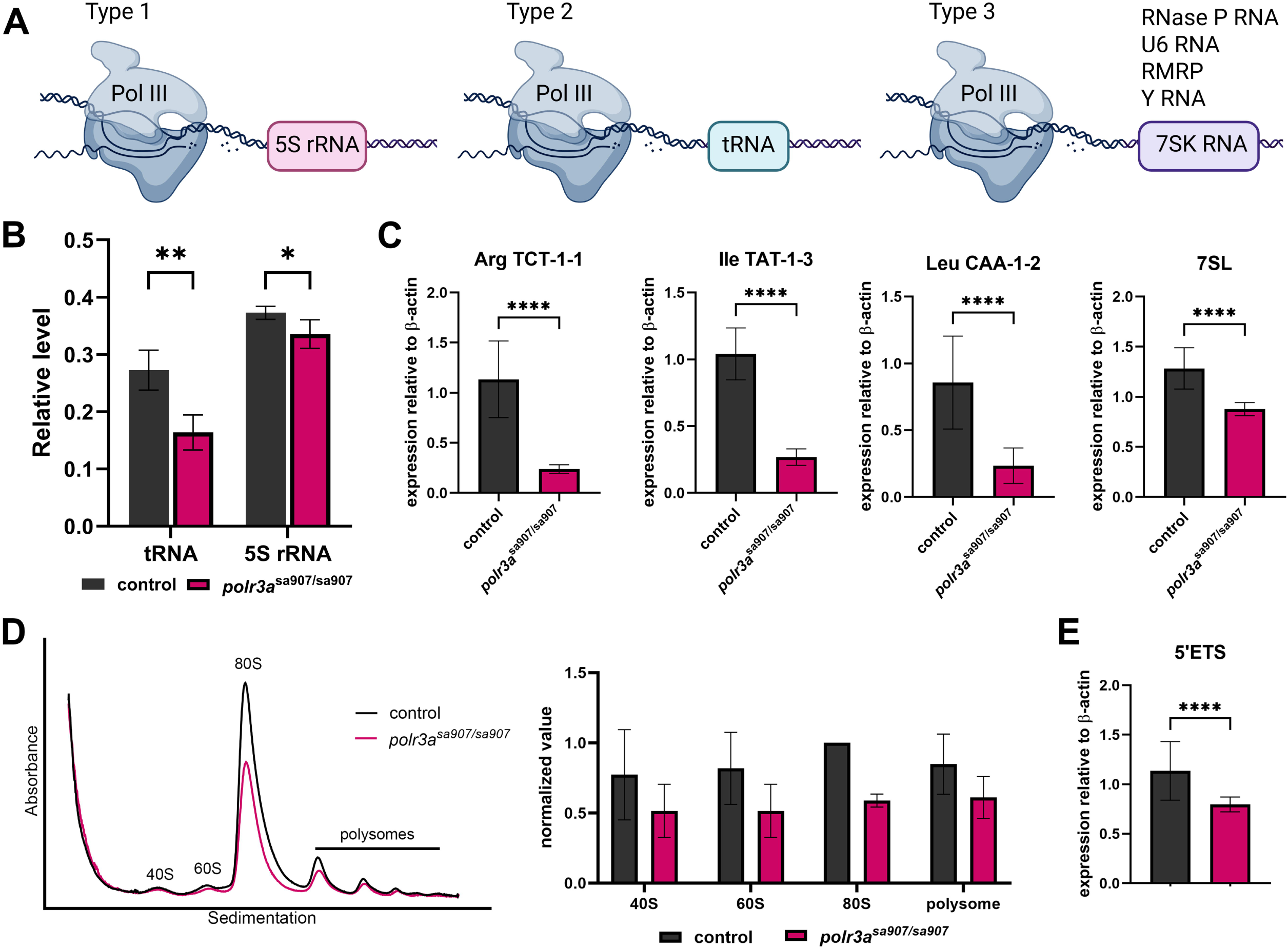
Pol III-mediated transcription and ribosome biogenesis are diminished in *polr3a^sa907/sa907^*zebrafish. (A) Pol III transcribes a variety of non-coding RNAs transcribed by three different types of promoters. Examples of genes transcribed by Pol III are represented. Created in BioRender. Watt, K. (2026) https://BioRender.com/qbgot2s (B) Analysis of total 5S rRNA and tRNAs on the Agilent bioanalyzer at 5 dpf shows significant reductions in *polr3a^sa907/sa907^* zebrafish. **p = 0.003; *p = 0.035. (C) qRT-PCR for pre-tRNAs and 7SL RNA at 3 dpf reveals consistent downregulation across all genes examined. ****p <0.0001. (D) Polysome profiling at 5 dpf reveals diminished ribosome assembly, showing significant reduction in the 80S peak and a trend of smaller polysome peaks. (E) At 3 dpf, *polr3a^sa907/sa907^* zebrafish show diminished 5’ETS expression, a region of the 47S rRNA transcribed by Pol I. ****p <0.0001.

### tRNA transcription and ribosome biogenesis are diminished in *polr3a^sa907/sa907^* zebrafish

Pol III transcribes tRNAs, rRNA, as well as a variety of non-coding RNAs that have essential cellular functions (Fig 6 A). Proliferating cells express a distinct set of tRNAs from those expressed in cells undergoing differentiation [56] and reduced ribosome biogenesis can trigger nucleolar stress and activation of p53 [58], resulting in increased cell death. We therefore hypothesized that diminished tRNA and rRNA transcription in *polr3a^sa907/sa907^* zebrafish contribute to the observed increased cell death and reduced proliferation. We assessed total 5S rRNA and tRNA levels at 5 dpf and found a significant reduction in both, relative to 5.8S rRNA (Fig 6 B). A greater reduction was observed in tRNA levels relative to the 5S rRNA, suggesting that not all genes transcribed by Pol III are equally affected in *polr3a^sa907/sa907^* zebrafish. To assess individual pre-tRNA levels, we performed qRT-PCR for intron-containing tRNAs at 3 dpf. This demonstrated a consistent reduction in expression level across all tRNAs examined, with an approximately 80% reduction in expression, and 7SL RNA levels were also significantly reduced by 40% (Fig 6 C). Together, this suggests that the *polr3a^sa907/sa907^* zebrafish display a reduction in Pol III-mediated transcription, especially for tRNAs.

As genes transcribed by Pol III have critical functions in ribosome biogenesis and translation, we assessed ribosome biogenesis using polysome profiling at 5 dpf. The 80S peak was reduced in *polr3a^sa907/sa907^* zebrafish, along with slight reductions in the polysome peaks, indicating reduced ribosome biogenesis and translation (Fig 6 D). The trend of reduction of all peaks in the polysome profile, not only the 60S where the 5S rRNA is incorporated, led us to hypothesize that Pol I-mediated transcription is also disrupted in *polr3a^sa907/sa907^* zebrafish. Reduced rRNA levels and nucleolar disruption have been observed in fibroblasts from individuals with WRS [59], raising the possibility of a feedback mechanism between Pol III and Pol I in *POLR3A*-associated disease. We assessed Pol I-mediated transcription via qRT-PCR for the 5’ETS region of the 47S rRNA and observed a 20% reduction in 5’ETS relative to controls at 3 dpf (Fig 6 E). Thus, Pol I-mediated transcription is reduced prior to the presence of obvious phenotypic changes in *polr3a^sa907/sa907^* zebrafish.

To understand when changes in rRNA transcription were occurring relative to changes in tRNA transcription in *polr3a^sa907/sa907^* zebrafish, we performed qRT-PCR at 2 dpf. These data show that pre-tRNAs are significantly downregulated by approximately 40% in mutants at 2 dpf while the 5’ETS region of rRNA remains unchanged, as does the 7SL RNA (S15 Fig). This demonstrates that Pol III-mediated transcription of tRNA is reduced prior to changes in rRNA transcription by Pol I. Together, this suggests that Pol III-mediated transcription is reduced in *polr3a^sa907/sa907^* zebrafish a full day prior to any phenotypic changes and that both Pol I and Pol III may contribute to reduced ribosome biogenesis and the observed craniofacial differences in *polr3a^sa907/sa907^* zebrafish after 3 dpf.

### Single cell RNA-sequencing reveals specific transcriptional changes in *polr3a^sa907/sa907^* zebrafish

To determine the molecular mechanisms by which mutation of *polr3a* affects craniofacial development, we performed single-cell RNA-sequencing (scRNA-seq). Whole heads from groups of control and mutant zebrafish were dissociated at 3 dpf to define transcriptional changes occurring prior to the onset of phenotypic differences. Over 30,000 cells per sample were sequenced to facilitate detection of rare cell types in the head, such as osteoblasts, and identify the most robust transcriptional changes between samples. After filtering, the data were normalized and low-resolution clustering (0.2) in Seurat resulted in 21 distinct clusters (Fig 7 A). Cell types were identified based on expression of cell-type specific markers, with cluster 11 identified as craniofacial cartilage cells based on high expression of *matn1, col2a1a,* and *sox9a* (S16 Fig). To determine global transcriptional changes in *polr3a^sa907/sa907^*zebrafish, we analyzed differentially expressed genes across all captured cells (Fig 7 B). Metascape analysis [60] of downregulated differentially expressed genes in *polr3a^sa907/sa907^* zebrafish revealed significant downregulation of processes including ribosomal large subunit biogenesis and regulation of the cell cycle (S17 Fig), consistent with our polysome profiling data (Fig 6 D) and analysis of pH3 (Fig 5). Metascape analysis of the upregulated differentially expressed genes in *polr3a^sa907/sa907^* zebrafish revealed significant enrichment of terms including amino acid metabolism and the p53 signaling pathway (S17 Fig).

**Fig 7.**
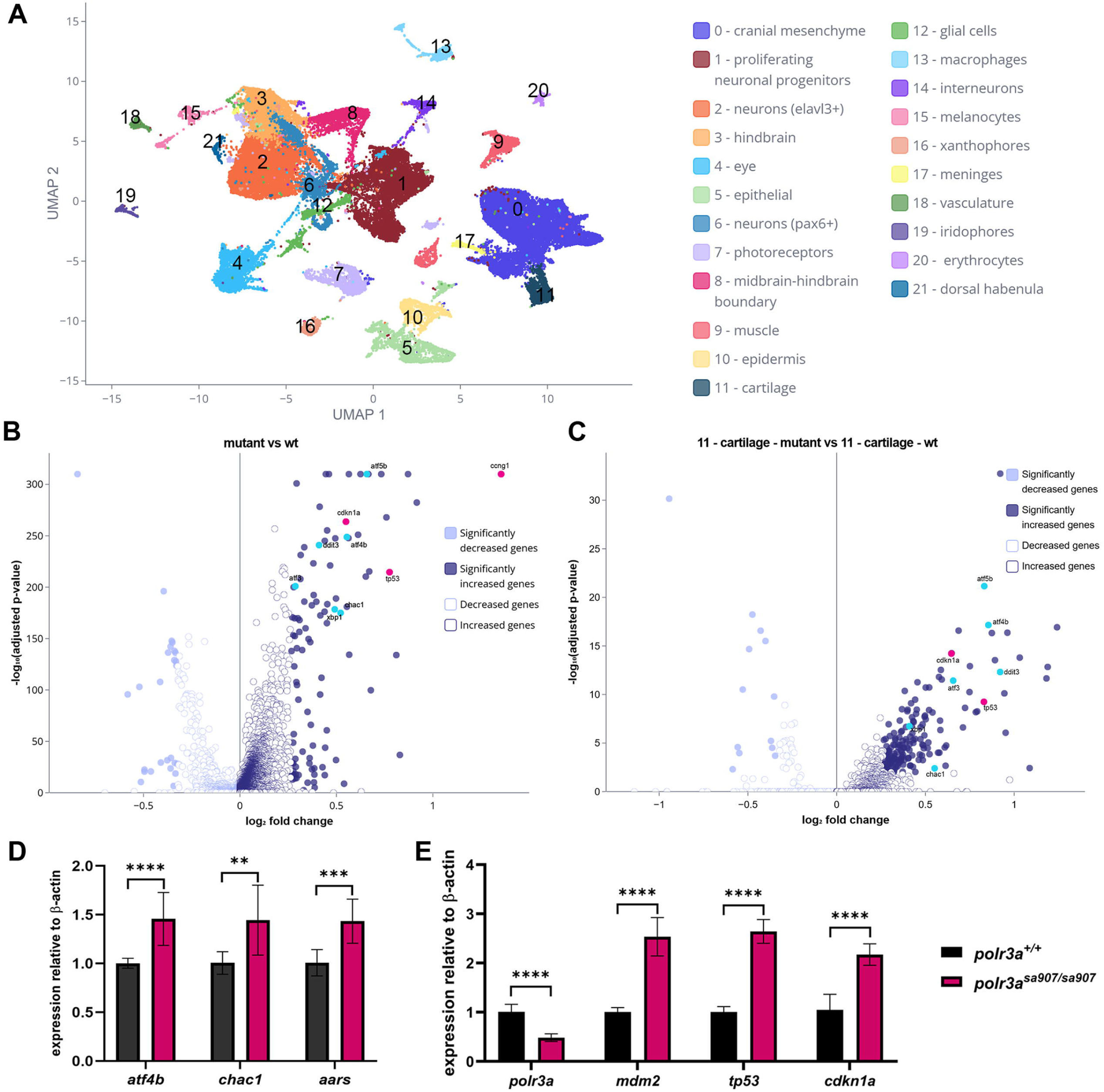
**Single cell RNA-sequencing reveals global transcriptional upregulation of genes in *polr3a^sa907/sa907^* zebrafish at 3 dpf**. (A) UMAP plot of aggregated mutant and wild-type cells. (B) Volcano plot of differentially expressed genes from a pseudo-bulk analysis of *polr3a^sa907/sa907^* cells compared to control cells. (C) Volcano plot of differentially expressed genes in *polr3a^sa907/sa907^* versus controls in the cartilage cluster. (D) qRT-PCR of genes in the integrated stress response pathway show about a 1.5-fold increase in mutants at 3 dpf. *atf4b* ****p <0.0001, *chac1* **p = 0.006, *aars* ***p = 0.0003. (E) qRT-PCR for genes in the Tp53 pathway (*mdm2*, *tp53*, and *cdkn1a*) show a nearly 3-fold increase in mutants at 3 dpf. ****p <0.0001.

Further investigation of the differentially expressed genes in these terms revealed upregulation of genes associated with the integrated stress response such as *atf4b, ddit3, chac1*, and *aars* (Fig 7 B, cyan). The integrated stress response (ISR) is a signaling network that converges on phosphorylation of eIF2α, resulting in activation of ATF4 and increased transcription of genes including *DDIT3* and *CHAC1* [61]. We also identified significant upregulation of genes in the p53 pathway including *tp53*, *mdm2*, and *cdkn1a* (Fig 7 B, magenta). Mdm2 is a negative regulator of Tp53, while *cdkn1a* is a transcriptional target of Tp53. To determine if these global changes in the ISR and Tp53 were occurring specifically in the developing craniofacial cartilage, we analyzed differentially expressed genes in cluster 11 (Fig 7 C).

Consistent with the global changes, we also observed upregulation of genes in the ISR pathway (Fig 7 C, cyan) and the Tp53 pathway (Fig 7 C, magenta). When we examined the expression across all cell types, we found that not all cells showed upregulation of these genes (S18, S19 Figs). This suggests that the loss of *polr3a* does not trigger the same response in all cells and that upregulation of both ISR pathway genes and Tp53 pathway genes occur in multiple cell types, including the craniofacial cartilage.

Next, to validate these changes, we performed qRT-PCR on RNA extracted from whole heads at 3 dpf. The relative expression of *atf4b, chac1,* and *aars* were upregulated in *polr3a^sa907/sa907^* zebrafish, with approximately a 1.5-fold increase in expression across these genes (Fig 7 D). The relative expression of Tp53 pathway genes including *mdm2, tp53*, and *cdkn1a* were all upregulated in *polr3a^sa907/sa907^* zebrafish, with approximately a 2.5 – 3-fold increase in expression relative to controls (Fig 7 E). Altogether, these data demonstrate the activation of multiple stress responses in *polr3a^sa907/sa907^*zebrafish, including increases in genes in the ISR pathway and Tp53 pathway genes, both broadly and within craniofacial tissues.

### *tp53* is increased in *polr3a^sa907/sa907^* zebrafish, but inhibition does not fully rescue cartilage and bone development

The Tp53 pathway showed significant upregulation in the craniofacial cartilage and Tp53 is a known regulator of both Pol I and Pol III [62–64]. We hypothesized that increased *tp53* could be mediating responses to reduced rRNA and tRNA and triggers cell death in *polr3a^sa907/sa907^* zebrafish, as disruptions in ribosome biogenesis are known to trigger upregulation of Tp53 [58]. Further, Tp53 activation has been observed in multiple models with disruptions in rRNA transcription, ribosomal proteins, as well as Pol III [21, 50, 51, 65–67]. We genetically inhibited *tp53* to test its contribution to craniofacial development in *polr3a^sa907/sa907^*zebrafish using the *tp53^M214K^* allele [68], referred to as *tp53^-/-^*. At 3 dpf, *polr3a^sa907/sa907^; tp53^-/-^* zebrafish show reduced cell death compared to *polr3a^sa907/sa907^*zebrafish (Fig 8 A-C; quantification in D), indicating that increased cell death at this stage is, at least in part, due to increased *tp53*. Interestingly, we did not observe any changes in global proliferation at this stage (S20 Fig). Given the reduction in cell death, we next examined the craniofacial cartilage and bone at 6 dpf in *polr3a^sa907/sa907^; tp53^-^* zebrafish and controls. Using Alcian blue and Alizarin red staining, we observed limited improvement in cartilage and bone at 6 dpf, despite the reduction in cell death at 3 dpf (S21 Fig). Although some improvement in the cartilage is evident in *polr3a^sa907/sa907^; tp53^-/-^* zebrafish compared to *polr3a^sa907/sa907^; tp53^+/+^* siblings, including slight increases in the length of the ceratohyal, these were not statistically significant and overall, the craniofacial cartilage elements and head length remain reduced in size in *polr3a^sa907/sa907^; tp53^-/-^* zebrafish (S21 Fig I-K). Similarly, we observe little improvement in bone staining regardless of *tp53* status (S21 Fig). In order to quantify the changes in overall bone size, we performed live Alizarin red staining and imaged the opercle at 6 dpf (Fig 8 E-G). Image analysis revealed that the morphology of the opercle improves slightly in *polr3a^sa907/sa907^; tp53^-/-^* zebrafish and there is a trend of an increase in overall volume compared to *polr3a^sa907/sa907^; tp53^+/+^* zebrafish. However, these improvements were not statistically significant and the opercle remains small relative to controls regardless of *tp53* status (Fig 8 H). To determine if bone development improved in *polr3a^sa907/sa907^; tp53^-/-^*zebrafish if given more time, we examined cartilage and bone at 8 dpf. Consistent with our observations at 6 dpf, this showed that while there are slight improvements in the size of cartilage elements, the branchiostegal rays and entopterygoid remain small or absent (S22 Fig). Further, *polr3a^sa907/sa907^; tp53^-/-^*zebrafish do not survive beyond 8 dpf, demonstrating that Tp53-independent mechanisms are important for overall survival.

**Fig 8.**
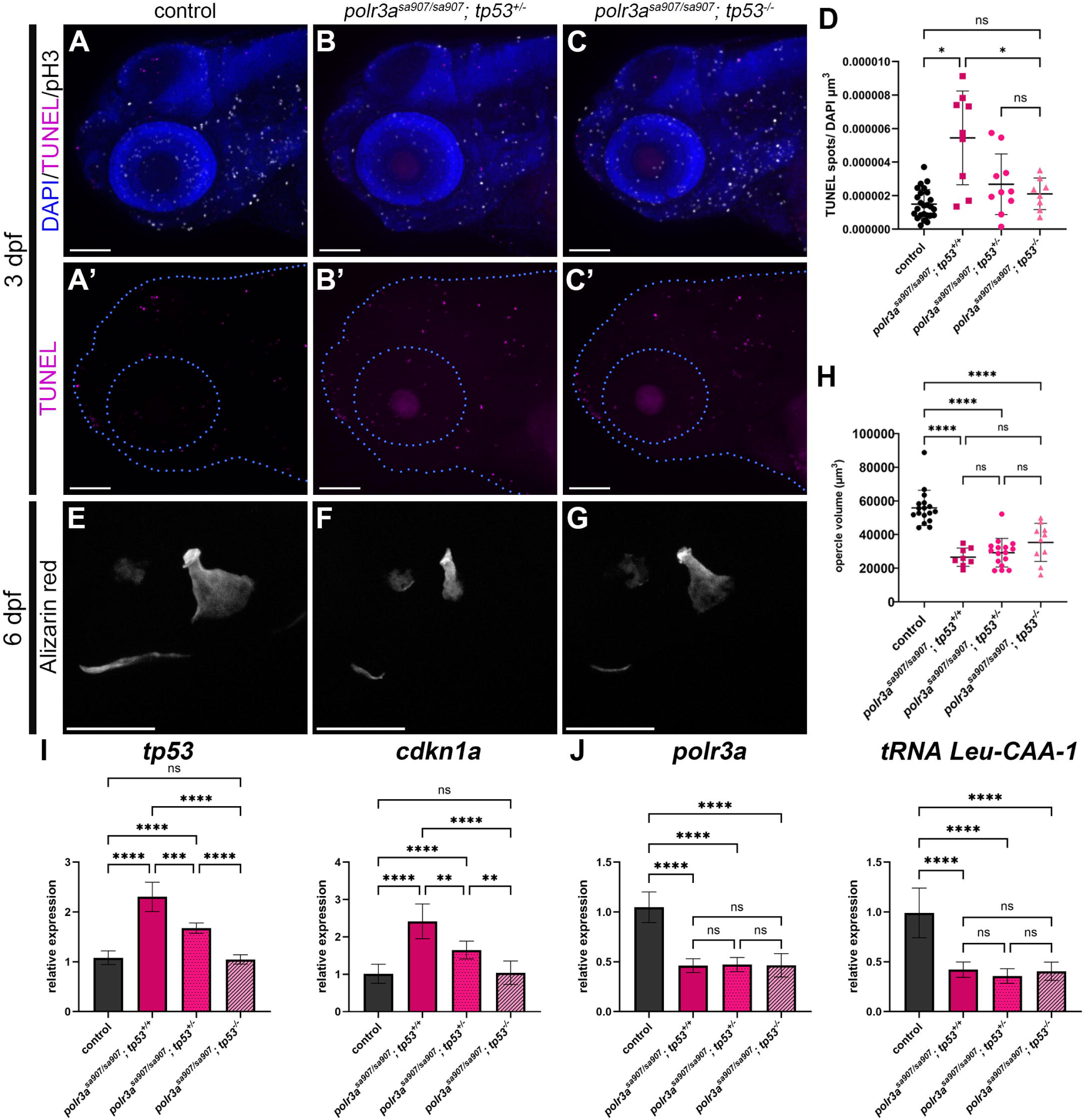
Genetic inhibition of *tp53* reduces cell death and leads to slight improvements in *polr3a^sa907/sa907^*zebrafish. (A-C) TUNEL (magenta) and pH3 (white) staining at 3 dpf. Quantification reveals a decrease in global cell death (G; *p = 0.013 control vs *polr3a^sa907/sa907^; tp53^+/+^*; *p = 0.038 *polr3a^sa907/sa907^; tp53^+/+^* vs. *polr3a^sa907/sa907^; tp53^-/-^*), but no change in proliferation (H; ns, not significant). n = 25 controls, n = 9 *polr3a^sa907/sa907^; tp53^+/+^,* n = 10 *polr3a^sa907/sa907^; tp53^+/-^,* n = 8 *polr3a^sa907/sa907^; tp53^-/-^.* (D-F) Alizarin red stain of the opercle at 6 dpf shows slight improvements in morphology, but no statistically significant improvement in the overall volume of the opercle in *polr3a^sa907/sa907^; tp53^-/-^* zebrafish (I; ****p <0.0001). n = 17 controls, n = 8 *polr3a^sa907/sa907^; tp53^+/+^,* n = 16 *polr3a^sa907/sa907^; tp53^+/-^,* n = 10 *polr3a^sa907/sa907^; tp53^-/-^.* Scale bar = 100 µm. (I) qRT-PCR for *tp53* and *cdkn1a* at 3 dpf demonstrates a dosage-dependent reduction in expression in *polr3a^sa907/sa907^* zebrafish based on *tp53* status. Expression levels of *tp53* and *cdkn1a* in *polr3a^sa907/sa907^; tp53^-/-^* zebrafish are not significantly different from controls. Adjusted p-values: **p < 0.005, ***p = 0.0007 ****p <0.0001. (J) qRT-PCR for *polr3a* and *tRNA-Leu-CAA-1-1* demonstrate that expression levels of these genes remain reduced in *polr3a^sa907/sa907^* zebrafish regardless of *tp53* status. Adjusted p-values: ****p <0.0001.

To test if there were transcriptional changes in *polr3a^sa907/sa907^; tp53^-/-^* zebrafish that precede these differences, we performed qRT-PCR at 3 dpf. This revealed a dose-dependent reduction in expression of *tp53* and *cdkn1a* in *polr3a^sa907/sa907^*zebrafish such that upon inhibition of both copies of *tp53*, there is no longer a significant difference between controls and mutants, consistent with the observed reduction in cell death (Fig 8 A-D). However, when we examined *polr3a* and one pre-tRNA gene, *Leu-CAA-1-1*, there were no differences in expression in mutants regardless of *tp53* status. This suggests that although *tp53* inhibition reduces cell death, there is not an improvement in Pol III-mediated transcription. Altogether, these data suggest that *tp53* is increased in *polr3a^sa907/sa907^*zebrafish and its inhibition reduces cell death at 3 dpf, and leads to slight improvements in the development of the craniofacial cartilage and bone. This data additionally suggests that Tp53-independent mechanisms also contribute to deficiencies in the development of the craniofacial skeleton in *polr3a^sa907/sa907^* zebrafish.

## Discussion

The tissue-specificity of developmental differences observed in ribosomopathies as well as Pol III-related syndromes remains an open question. The presence of craniofacial differences associated with Pol III indicates that its function is critical for normal craniofacial development. However, our understanding of how differences in Pol III-mediated transcription affect NCC and craniofacial development and the underlying mechanisms remains incomplete. In this study, we demonstrated that Pol III subunit *polr3a* is required for craniofacial development in zebrafish through regulation of tRNAs and ribosome biogenesis which influence cell survival (Fig 9).

**Fig 9.**
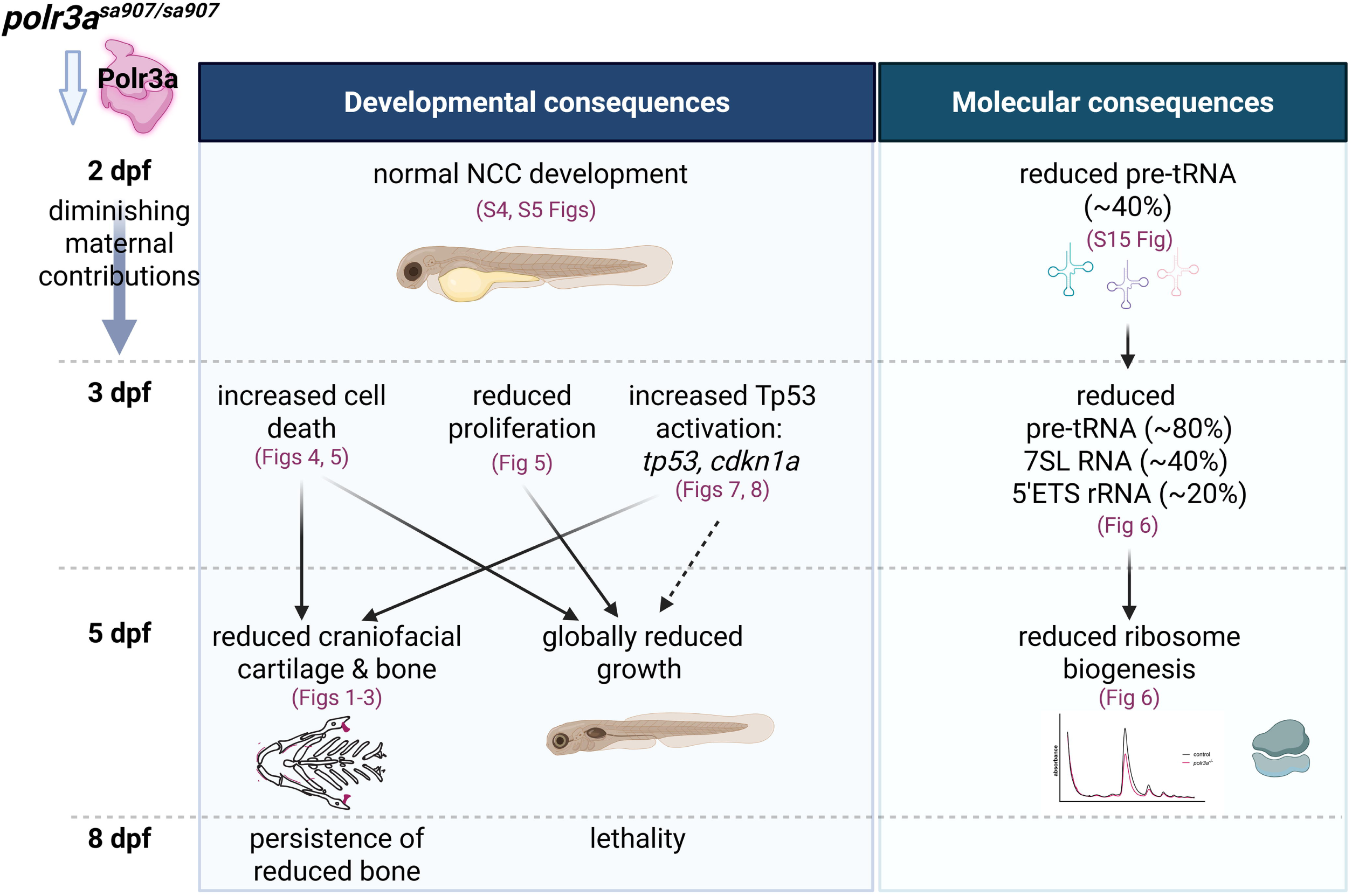
Summary of the consequences of *polr3a^sa907^* in zebrafish development. At 2 dpf, no morphological differences are apparent in *polr3a^sa907/sa907^*zebrafish and neural crest cells (NCC) develop normally. However, at this same stage, slight reductions are observed in pre-tRNA transcription. By 3 dpf, after depletion of maternal stores, *polr3a^sa907/sa907^* zebrafish show increased cell death and reduced proliferation, which contributes to a global reduction in growth. Increased cell death is also observed in the developing craniofacial cartilage and scRNA-seq revealed elevated expression of the *tp53* pathway in the cartilage. Increased *tp53* is also observed in some other cell types, contributing to overall cranial development (dashed arrow). On the molecular side, by 3 dpf, transcription of pre-tRNAs are significantly reduced, as are other genes transcribed by Pol III including 7SL RNA. A slight reduction in 5’ETS rRNA transcribed by Pol I is also observed. Combined, this leads to an overall reduction in ribosome biogenesis at 5 dpf. Developmentally, at 5 dpf, there are measurable differences in the craniofacial cartilage and bone, with reductions in all elements examined. These differences persist through 8 dpf, which is the limit of the lifespan of *polr3a^sa907/sa907^*zebrafish. Created in BioRender. Watt, K. (2026) https://BioRender.com/ni7bxps

One frequently proposed mechanism for the tissue-specificity of ribosomopathies has been that perturbations in ribosome biogenesis affect highly proliferative cell types, such as NCCs, more than others. Further, the frequency of craniofacial differences in ribosomopathies has led to the hypothesis that NCCs are especially sensitive to disruptions in ribosome biogenesis. For example, craniofacial differences are defining diagnostic criteria for ribosomopathies such as Treacher Collins syndrome [20, 69–71]. Animal models for Treacher Collins syndrome showed increased neuroepithelial cell death of NCC progenitors [21, 30, 72], with reduced craniofacial cartilage and bone by 5 dpf in zebrafish [21]. Studies further demonstrated that NCCs have a relatively high requirement for ribosome biogenesis [30]. The *polr3a^sa907/sa907^*zebrafish showed reduced craniofacial cartilage and bone by 5 dpf (Fig 1), which led us to hypothesize that NCC development was disrupted. Surprisingly, we did not detect changes in NCC formation or migration (Fig 2; S4 Fig; Fig 9), in contrast to differences in the NCC within the pharyngeal arches by 36 hpf in *polr1c* and *polr1d* mutant zebrafish [21]. Further, no differences were identified in early differentiation into craniofacial cartilage and bone in *polr3a* mutants (S7 Fig, Fig 3). Increased cell death and reduced proliferation have been observed in affected tissues of other Pol III mutant models [14, 65, 73], and we observe global changes in cell death and proliferation in *polr3a* mutants (Fig 5).

Intriguingly, we did not identify changes in proliferation within the cartilage (Fig 4), which suggests a proliferation-independent mechanism in this tissue. Thus, our data suggest that there are additional mechanisms beyond proliferative capacity which contribute to phenotypic outcomes.

The diversity of affected tissues in ribosomopathies and the observation that mutations in Pol III subunits result in distinct phenotypes in animal models also suggest that there are factors beyond proliferative capacity that influence phenotypic outcomes. Conditional mutations in *Polr3a* at postnatal stages in mice result in changes in cerebral cortex and the exocrine pancreas, while other tissues are unaffected [74]. Mutations in Pol III subunit *polr3b* affect pancreas development in zebrafish [73], while mutations in the Pol III subunit *rpc9 (crcp)* affect hematopoietic development [65]. Craniofacial development in these models was not assessed, so it is possible that there are also differences in cranioskeletal development in addition to these other tissues. Furthermore, in humans, the genotype-phenotype correlations of many variants in Pol III are not well understood. Differences in phenotypic presentations may be due to a combination of the specific variants, differences in genetic background, tissue-specific sensitivities to deficient translation, and other factors. Together, these data raise the possibility that there are different requirements for Pol III across tissues and that the degree of disruption to Pol III could influence these outcomes.

To better understand different requirements for Pol III across developmental stages, we examined mRNA expression levels. We detected high maternal expression of *polr3a* (S11 Fig), but how long maternal tRNAs persist during zebrafish embryogenesis is unknown. Studies in embryonic chick muscle have suggested the half-life of tRNAs is around 50 hours [75], which would be consistent with the lack of observed phenotypes in Pol III mutant zebrafish until after 2 dpf. Further, no developmental differences were noted prior to 2 dpf in other Pol III mutant zebrafish models [65, 73], while morpholino-mediated knockdown of *polr3b* results in lethality prior to 24 hpf [73]. In contrast, differences in craniofacial development are apparent as early as 24 hpf in *polr1c* and *polr1d* mutant zebrafish [21], much earlier than *polr3a^sa907/sa907^*zebrafish. Altogether, this suggests that *polr3a* has distinct requirements during craniofacial development and that extensive maternal contribution of Pol III and tRNAs may support development for the first two days in zebrafish.

Pol III transcribes many noncoding RNAs which have functions in ribosome biogenesis, transcription, and translation, including tRNA, 5S rRNA, RMRP, 7SK RNA, and 7SL RNA. These RNAs are critical for cell survival and proliferation in multiple tissues and structures. Diminished tRNA transcription was found in multiple Pol III models including mice [74], zebrafish [65, 73], and human cells [76–78], although there were differences in which tRNAs showed downregulation and how tRNAs were assessed. Consistent with these studies, we identified reduced transcription of tRNAs in *polr3a^sa907/sa907^* zebrafish (Fig 6 B,C), and this was the first change we could detect at 2 dpf (S15 Fig). Tissue-specific expression of tRNAs has been noted in multiple model systems [56, 79], leading to the hypothesis that changes in tRNA levels may affect some tissues more than others. In contrast to this, recent studies have suggested that, while tRNA pools do change during differentiation, these tRNA pools remain buffered at the level of the anticodon [80]. In zebrafish, a switch in global translation, and not changes to the tRNA pool, is important at the zebrafish maternal to zygotic transition [81] and changes in tRNA abundance do not strongly correlate with changes in codon frequency at multiple stages of zebrafish development [82]. These studies do not rule out that changes in the tRNA pool can have specific effects, but raise the possibility that additional factors or perhaps the overall transcriptional and translational requirements of different tissues contribute to the specificity of Pol III-related syndromes.

Intriguingly, we observed diminished transcription of rRNA in *polr3a^sa907/sa907^*zebrafish (Fig 6 E), as well as a trend of diminished ribosome biogenesis and translation (Fig 6 D), suggesting that reduced Pol III activity also influences Pol I and ribosome biogenesis. Reduced rRNA levels and nucleolar disruption were observed in *POLR3A*-associated Wiedemann-Rautenstrauch syndrome fibroblasts [59] and conditional deletion of *Brf1* in mice led to diminished ribosome biogenesis [8], further supporting the idea that there is feedback from perturbation of Pol III on ribosome biogenesis. In yeast *rpc160Δ* mutants, the *polr3a* homolog, rRNA levels were reduced upon reduced transcription of tRNAs such that the tRNA:rRNA ratio was restored [83]. This raises the possibility that Pol III-related syndromes could either directly affect ribosome biogenesis through 5S rRNA or RMRP levels, or indirectly through feedback from reduced tRNA levels. Diminished tRNAs could lead to reduced translation of ribosomal protein genes, further affecting ribosome biogenesis. From the scRNA-seq data, we observed reduced expression of some ribosomal protein genes in *polr3a^sa907/sa907^* zebrafish, and therefore this could be an additional mechanism leading to reduced ribosome biogenesis that would be worth investigating in future studies. The combined reduction in Pol I and Pol III, as well as ribosome biogenesis, could have broad effects on cell survival, but how this results in tissue-specificity remains unclear.

Our scRNA-seq revealed that there are multiple transcriptional responses in *polr3a^sa907/sa907^* zebrafish, including activation of both the ISR and Tp53 (Fig 7; S16-S19 Figs). These pathways were identified within cartilage cells as well as other cell types, demonstrating that some stress responses are activated across multiple cell types. Activation of the ISR has been reported in other models with disruptions in tRNA biogenesis [74, 84–86], suggesting that upregulation of genes in the ISR pathway may be in response to diminished transcription of tRNAs by Pol III. ISR activation has not been investigated in either *polr3b* or *crcp* mutant zebrafish models, so it remains unclear if this is a common response to the loss of Pol III across models and tissues. Our data demonstrate that activation of the ISR is not unique to neuronal cell types or brain, which have been the focus of prior studies [74, 85], and appears to occur in nearly all cells in *polr3a^sa907/sa907^* zebrafish at 3 dpf (S18 Fig).

Previous studies have shown that the degree and timing of Tp53 activation can lead to diverse phenotypic outcomes [87]. Consistent with this model, increased expression of *tp53* appeared to be more limited to craniofacial tissues and some neuronal cells in *polr3a^sa907/sa907^*zebrafish (pathway upregulation in 3/22 clusters; S19 Fig). Inhibition of *tp53* improved overall cell survival at 3 dpf in *polr3a^sa907/sa907^* zebrafish (Fig 8) and led to subtle improvements in cartilage and bone development (Fig 8; S21, S22 Figs), demonstrating that some cell survival is regulated by Tp53. However, *tp53* inhibition did not improve the overall survival of *polr3a^sa907/sa907^*zebrafish. In contrast, *tp53* inhibition in the *crcp* mutant zebrafish completely rescued hematopoietic development [65]. This suggests that *tp53* upregulation is not the only reason for diminished survival and growth of the craniofacial cartilage and bone in *polr3a* mutants and that there must be additional contributing factors. One possibility is that transcription by Pol III is required for osteoblast differentiation, which has been shown in cell culture models [88], and that the tRNA supply is simply insufficient for the demands of differentiation. In support of this, we did not observe an improvement in pre-tRNA transcription upon *tp53* inhibition in *polr3a* mutants (Fig 8 J), although it is important to note that our analysis was limited in scope and there could be changes that we did not detect. Given the additional phenotypes present in our *polr3a* mutant zebrafish model, this tRNA-dependent mechanism could be affecting differentiation of additional cell types beyond osteoblasts. It is also possible that the limited rescue of cartilage and bone development upon Tp53 inhibition is due to contributions from alternative pathways, such as the ISR. If activation of the ISR halts translation independently of Tp53, then it is possible that some of the craniofacial cartilage and bone hypoplasia is mediated through this pathway.

Altogether, our data demonstrate a role for Pol III in craniofacial development and provide new evidence that multiple stress responses are activated upon the loss of Pol III. Different responses, such as Tp53 activation, are not ubiquitously activated. This raises the intriguing possibility that the type and degree of stress response may also contribute to tissue-specificity of developmental differences in POLR3-related syndromes. Additional studies are needed to determine the tissue-specific mechanisms regulating craniofacial growth and differentiation and whether different mutations result in differences in Pol III-mediated transcription which ultimately influence cell fate.

## Study Limitations

In this study, we utilized a zebrafish model to understand the role for *polr3a* in craniofacial development. As zebrafish have maternal contribution of ribosomes and *polr3a*, our model is not a full null. We hypothesize that maternal contributions result in the relatively late onset of developmental differences in *polr3a^sa907/sa907^* zebrafish and that the timing of depletion of maternal deposits contributes to the observed phenotypes. However, if this is the reason for the relatively mild phenotypes observed, all tissues that develop at later stages would be predicted to be affected after the maternal contribution is depleted. In contrast to this prediction, *mbp:tagRFP* expression was unchanged in *polr3a* mutants (S12 Fig). As pathogenic variants in *POLR3A* are associated with POLR3-HLD, the lack of a difference in *mbp* expression was surprising. It is possible we did not detect changes because of our limited analysis and that the production of myelin, which we did not directly assess, could be affected. Alternatively, there might be differences observed at later developmental stages if the zebrafish survived past 8 dpf. Thus, the phenotypes we observe are reflective of a reduction in *polr3a*, but the developmental timing and how long maternal contributions last must be considered in the interpretation of phenotypes. Further, in our study we focused on understanding craniofacial cartilage and bone, but additional tissues and structures are affected in *polr3a^sa907/sa907^* zebrafish. Additional examination of these tissues is necessary to determine the full phenotypic presentation of *polr3a* mutant fish and the degree to which additional tissues are affected.

## Materials and methods

### Zebrafish maintenance and husbandry

All zebrafish work was approved by the Institutional Animal Care and Use Committee at the University of Colorado Anschutz Medical Campus (Protocol# 01361). Zebrafish (*Danio rerio*) were maintained on an AB background and housed at 28.5°C on a 14/10 hour light/dark cycle. Adult zebrafish stocks were maintained as heterozygotes. Heterozygous *polr3a^sa907^* fish, obtained from ZIRC (ZL10320.03), were crossed with additional lines including *Tg(-7.2kb-sox10:egfp)* ([35], gift from T. Schilling lab), *Tg(sp7:EGFP)^b1212^*([41], gift from J. Nichols lab), *sox9a^zc81Tg^* referred to as *sox9a:egfp* ([37], gift from J. Nichols lab), *Tg(mbp:tagRFP)* ([53], gift from B. Appel lab) and *tp53^M214K^* ([68], ZIRC stock zdf1). Embryos were raised in 0.5x E2 media at 28.5°C and staged according to [89]. When necessary, 1-phenyl-2-thiourea (0.003%) was added to embryo media to prevent pigment development.

### Genotyping

*polr3a^sa907^* zebrafish samples were genotyped either by PCR followed by Sanger sequencing or by KASP assay. For samples genotyped by PCR and Sanger Sequencing, DNA was extracted and a 468 bp PCR product spanning the sa907 locus was amplified using F: 5’- GGCTCTTCCTCCTGAAGTTATT-3’ and R 5’-AAAGGCAAAAGGGGTCCAAC-3’. The PCR product was cleaned using ExoSAP-IT reagent (Fisher, 750011EA) and sent for Sanger sequencing using primer 5’-CTGCTGTCTTTGTGTGGAGG-3’.

Sequences were analyzed for the presence of ‘C’ or ‘T’ at the sa907 locus. For samples genotyped by KASP assay, KASP master mix with High Rox (LGC) and KASP assay mix 3541.02 (LGC) were used to prepare the genotyping reaction according to the manufacturer’s protocol. Samples were amplified on an Applied Biosystems StepOnePlus Real-Time PCR System and analyzed. PCR was used to genotype the *tp53^M214K^* allele. Wildtype primers were F: 5’ AGCTGCATGGGGGGGAT-3’ and R: 5’-GATAGCCTAGTGCGAGCACACTCTT-3’ and mutant primers were F: 5’ AGCTGCATGGGGGGGAA-3’ and R: 5’- GATAGCCTAGTGCGAGCACACTCTT-3’.

### Skeletal staining

Alcian blue and Alizarin red staining was performed on samples as previously described [90]. In brief, larvae were collected at 5, 6, or 8 dpf and fixed in 2% PFA at room temperature for 1 hour. Larvae were then stained overnight with 0.04% Alcian blue, rehydrated using a graded EtOH series, and stained with 0.01% Alizarin red. Larvae were equilibrated in 75% glycerol/0.1% KOH and prepared for imaging.

Images were captured on a Leica M205 FCA stereoscope equipped with a Leica DMC6200 Digital Camera using the Leica Application Suite X software (version 5.1.0.25446). Manual z-stacks were acquired and maximum projection images were generated using Helicon Focus software (version 8.3.0). Images were quantified using ImageJ using the markings indicated in S2 Fig. Specifically, for the length of the ceratohyal, both left and right sides were measured and then the average of the two was used for statistical comparison. For the width of Meckel’s, the distance from jaw joint to jaw joint was measured; head length was measured from the anterior most point of the ethmoid plate to the limit of Alizarin staining in the notochord; and eye diameter was measured. To account for differences in imaging across multiple clutches, normalized values were used in S21 Fig. For these calculations, the average of all control measurements was used to normalize values within a clutch. For each clutch, each measurement was divided by the average control. After calculating these normalized values, measurements were pooled from three different clutches and then used for statistical comparisons. For live Alizarin red staining, larvae were stained with 0.01% Alizarin red diluted in E2 media for 20 minutes and then rinsed several times with E2 media. Larvae were anesthetized in MS-222 and mounted in 0.8% low melting point agarose containing MS-222 and imaged on a Leica DMi8 microscope equipped with an Andor Dragonfly 301 spinning disk confocal system.

### In Situ Hybridization, immunostaining, and TUNEL

#### In situ hybridization

DIG-labeled *col2a1a* probe was generated from *col2a1a* plasmid digested with HindIII (NEB; R0104M) and in vitro transcribed with T3 RNA polymerase (NEB; M0378S) and DIG nucleotide labeling mix (Roche, #11277073910). RNA was precipitated and reconstituted in 50% formamide. In situ hybridization was completed according to standard protocols. Briefly, embryos were permeabilized with Proteinase K and hybridized in probes diluted to 2.5 ng/μl in hybridization buffer (50% formamide, 5X SSC pH4.5, 1% SDS) overnight at 68°C. The probe was removed and embryos were washed with TBS + Tween20 and blocked for a minimum of 1 hour with blocking reagent (Roche, Catalog# 11096176001) prior to incubation in Anti-Digoxigenin-AP, Fab fragments (1:2000; Roche, Catalog# 11093274910). Signal was detected using NBT/BCIP and the development reaction was stopped when deemed complete. Embryos were rinsed with TBS and stored in 4% PFA to stop the reaction and then imaged with the Leica M205 FCA stereoscope described above.

##### *<u>In situ Hybridization Chain Reaction (HCR)</u>* for c*dh16* was conducted following Molecular Instruments

Whole Mount Zebrafish protocol (https://files.molecularinstruments.com/MI-Protocol-RNAFISH-Zebrafish-Rev10.pdf). HCR for *polr3a* was conducted with slight modifications. Samples were incubated for 3 days in 20 nM of probe solution at 37°C. Samples were mounted in 0.8% LMP agarose in PBS and imaged on Leica DMi8 microscope equipped with an Andor Dragonfly 301 spinning disk confocal system using the 10x objective.

*Immunostaining* was performed as previously described with some modifications (Westerfield 2000).

Embryos and larvae were fixed overnight in 4% PFA and then dehydrated using a graded MeOH/PBT series and stored at least overnight at –20°C. Samples were rehydrated using a graded MeOH/PBT series and digested with Proteinase K (Invitrogen, 1:2000). Primary antibodies used include anti-GFP (Invitrogen A66455, 1:500; RRID:AB_221570), anti-pH3 (Millipore-Sigma 05-806, 1:2000; RRID:AB_310016), and Alexa 546 conjugated phalloidin (Invitrogen, A22283). Secondary antibodies used include Alexa-488, Alexa-546 and Alexa-641 (Invitrogen, 1:500). DAPI staining was performed using a working concentration of either 2 ng/ul or 10 ng/ul diluted in PBS.

*<u>TUNEL</u>* staining was performed using the Roche TUNEL kit (Millipore-Sigma 12156792910) according to the manufacturer’s protocol with slight modifications. Samples were incubated in labeling solution for 1 hour on ice followed by a 1-hour incubation at 37°C. Samples were washed and then prepared for imaging.

### Confocal Imaging and quantification

*<u>Confocal Imaging</u>*: Fixed samples were mounted in 0.8% LMP agarose in PBS and imaged on a 3I Marianas inverted spinning disc confocal microscope using Slidebook software (version Slidebook6 2025). Whole head images were taken using a 10x objective, while pharyngeal arch 1 and 2 derived structures were imaged at 20x. Max projection images were either generated in the Slidebook software or ImageJ and scale bars were added in ImageJ. *sp7:gfp* and *mbp:tagRFP* zebrafish were mounted in 0.8% LMP agarose in E2 media with MS-222, while *sox9a:egfp* zebrafish were fixed in 4% PFA for 1 hour at room temperature and then mounted in 0.8% LMP agarose. Zebrafish were imaged on a Leica DMi8 microscope equipped with an Andor Dragonfly 301 spinning disk confocal system at 10x, 20x, or 40x. Max projection images were generated in ImageJ.

*<u>Image Quantification:</u>* Confocal images were analyzed using Imaris software (version 9.2.0). For analysis of global proliferation and cell death, a region of interest (ROI) was generated to include the whole head using the DAPI channel and all other channels were masked for the ROI. Spots tool was used to quantify the number of TUNEL positive or pH3 positive spots within the ROI. To analyze pharyngeal arch 1 and 2 derived structures in *sox10:egfp* zebrafish, the ROI was generated using *sox10:egfp* and then all other channels were masked to this ROI. TUNEL and pH3 were quantified using the spots tool. Automatic thresholding in Imaris software was applied to all samples. For quantifications of the opercle, cranial ganglia, and myelin basic protein, a ROI was set around the structure of interest. For quantifications of *sp7:gfp* and Alizarin red, the ROI was defined around the boundaries of the bone element of interest (opercle or dentary) based on *sp7:gfp* expression and the same ROI was used for both *sp7:gfp* and Alizarin red. For quantification of *mbp:tagRFP*, the ROI was defined as 1000 pixels in width. Threshold settings were the same for all animals from the same clutch. Volume was recorded for each sample. For *sox9a:egfp* quantifications, a ROI was created around Meckel’s, including the jaw joint. The ROI was then masked, and the spots tool was used to count cells within the masked volume. A manual threshold was selected and applied to all samples from the same clutch. Volume and spot number were recorded for each sample.

*<u>CellProfiler Analysis</u>*: *sox9a:egfp* positive cells were analyzed using Fiji (version 2.16.0/1.54p) and CellProfiler (version 4.2.6). Meckel’s cartilage was digitally isolated in Fiji using the polygon selection tool. A pipeline was developed in CellProfiler to quantify *sox9a:egfp* positive cell count, cell area, and cell eccentricity. The CellProfiler pipeline converted the images to grayscale using ColorToGray, identified *sox9a:egfp* positive cells using IdentifyPrimaryObjects, measured cell count, cell area, and cell eccentricity using MeasureObjectSizeShape, and exported the data to Microsoft Excel. IdentifyPrimaryObjects used a three-class global Otsu thresholding strategy with a smoothing scale of 1.3488 (Gaussian sigma of 1), threshold correction factor of 1, threshold bounds of 0.0-1.0, allowed object diameter of 10-75 pixels, and distinguished between clumped objects using intensity. Total *sox9a:egfp* positive cell counts were recorded for each larvae and graphed in GraphPad Prism (version 10.6.1). For each larvae, *sox9a:egfp* positive cell area was recorded for each *sox9a:egfp* positive cell and averaged on a per larvae basis. Similarly, for each larvae, *sox9a:egfp* positive cell eccentricity was recorded for each *sox9a:egfp* positive cell and averaged on a per larvae basis. Average *sox9a:egfp* positive cell area per larvae and average *sox9a:egfp* positive cell eccentricity per larvae were graphed in GraphPad Prism.

*<u>Image adjustments:</u>* Figures were assembled in Adobe Photoshop. Any adjustments to brightness or contrast were applied equally across the entire image.

### Sensorimotor Behavior Testing and Analyses

Larvae were relocated to the behavior room 30 minutes prior to testing and acclimated in an incubator at 28°C. To test acoustic startle response thresholds, each larva was placed into an individual well on a 36-well plate. Larvae were exposed to six sets of acoustic stimuli of increasing intensity as previously described [46, 91]. Stimulus intensity was measured with an accelerometer (PCB Piezotronics) and is reported in *g* (acceleration due to gravity). Stimulus intensities used range from 0.5370*g* to 51.1770*g*.

Each stimulus was administered five times and stimuli were separated by a 40-second inter-stimulus interval. After measuring acoustic startle thresholds, habituation was measured by presenting 40 stimuli with a three-second inter-stimulus interval. To capture larval movements, a high-speed camera (FASTCAM Mini UX50 Type 160-M-32G) was used to record at 1000 frames per second. For analysis, behavior videos were background-subtracted and tracking was performed using Flote [92, 93]. Behavior calls were made using Batchan and GraphPad Prism was used to perform statistical analysis.

### RNA extraction and qRT-PCR

RNA was collected from AB whole embryos or *polr3a* embryo heads at 2 – 5 dpf. For *polr3a* and *polr3a; tp53* RNA, heads were cut at the pectoral fin and stored in RNA later while tails were lysed and genotyped. Heads were then grouped based on genotype and RNA was extracted using Trizol RNA extraction according to the manufacturer’s protocol. RNA was cleaned up using either the Qiagen miRNeasy micro kit (Cat. # 217084) with on-column DNase treatment (Cat. # 79254) or the ZYMO RNA clean and concentrator kit (ZYMO, R1013) with DNase treatment. RNA concentration was checked on the Nanodrop 1000. RNA input was normalized for the cDNA synthesis reaction. cDNA was synthesized using the SuperScript IV First Strand Synthesis Kit (Invitrogen 18091050). Assay validations were conducted on wild-type samples for all primer pairs (See Table 1) to determine appropriate cDNA concentrations for amplification. Master mixes were prepared with iTaq Universal SYBR Green Supermix (Bio-Rad 1725121) and cDNA from 3 biological replicates with at least two technical replicates were pipetted into 384-well plates using the epMotion 5073 liquid handler. Plates were run on a BioRad CFX384 Real-Time PCR detection instrument at an annealing temperature of 60°C. Data was analyzed using the 2^-ΔΔCT^ method.

**Table 1.**
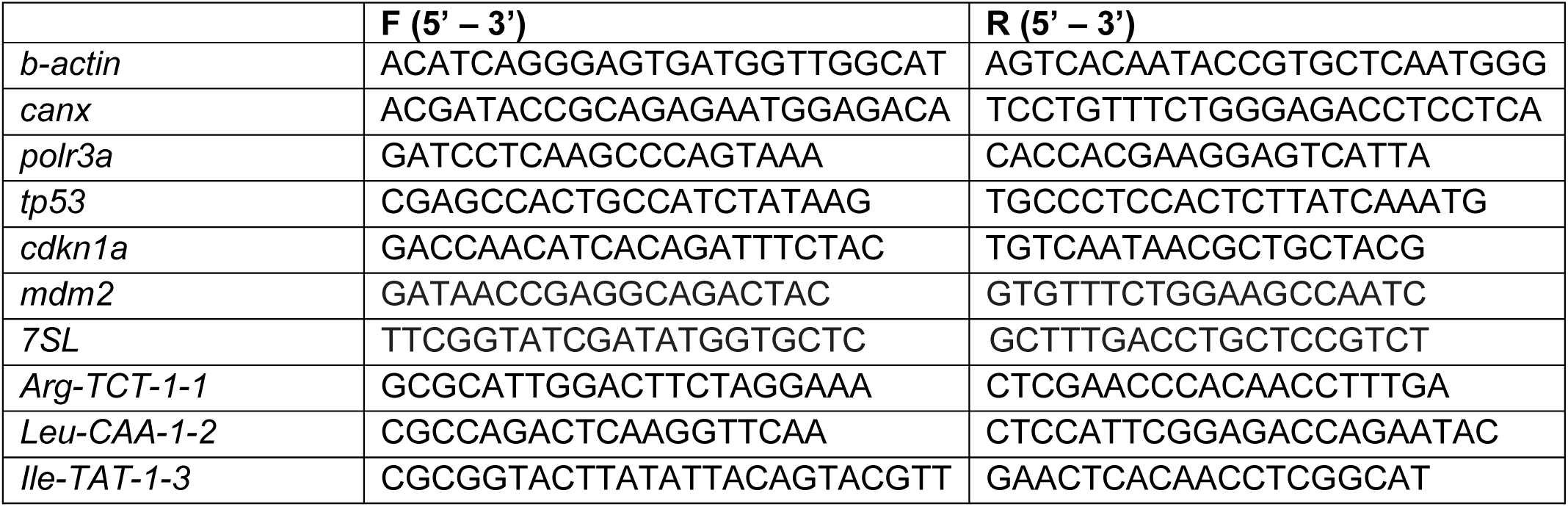
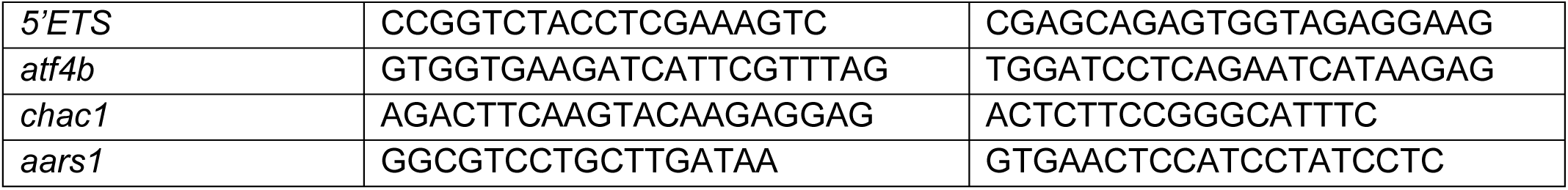
qRT-PCR primers.

### Bioanalyzer measurements

RNA was extracted from 5 dpf *polr3a* control and mutant zebrafish, as above. One microliter of RNA was loaded onto a small RNA chip for 5 mutant and 5 control samples and read on the Agilent 2100 Bioanalyzer. The area under the curve of individual peaks was determined in ImageJ and the 5S rRNA and tRNA peaks were normalized to the 5.8S peak as in [73]. Normalized values were used for statistical comparison between control and mutant samples.

### Western Blot

Protein was extracted from groups of genotyped zebrafish at 5 dpf in protein lysis buffer (50 mM Tris pH 8, 150 mM NaCl, 10% glycerol, 1% NP-40, 2 mM EDTA, 1X protease inhibitor, 1 mM PMSF, 10 mM NaF, 1 mM Na_3_VO_4_, 25 mM β-glycerophosphate). Lysates were cleared by centrifuging for 20 minutes at 13,000 x g. Protein samples were separated by SDS-PAGE gel, transferred to a PVDF membrane, and then blotting was performed according to standard protocols with HRP-conjugated secondary antibodies. Primary antibodies used were Polr3a (Invitrogen, PA5-21730; RRID:AB_11152880; 1:2000) and α-tubulin, (Sigma, T9026, RRID:AB_477593; 1:10000). Following blotting for Polr3a, blots were stripped with stripping buffer (2% SDS, 62.5 mM Tris HCl pH 6.8, 8 μL/mL of β-mercaptoethanol) for 30 min at 50°C, washed with TBS+Tween, and re-probed for α-tubulin as a loading control. Blots were developed using Pierce ECL Western Blotting Substrate (ThermoFisher) and imaged using a Bio-Rad ChemiDoc XRS+. Band intestities were analyzed in ImageJ and Polr3a values were normalized to α-tubulin loading controls.

### Polysome profiling

Zebrafish were sorted by phenotype at 5 dpf and 150 animals were used for each biological replicate. Polysome profiling was conducted as previously described [50] with slight modifications. Zebrafish were anesthetized in MS-222, rinsed with ice cold PBS with cycloheximide, and then dissociated in ice-cold lysis buffer (10 mM Tris-HCl (pH 7.4), 5 mM MgCl_2_, 100 mM KCl, 1% Triton X-100, 2 mM DTT, 500 U/ml RNasin, 100 μg/ml cycloheximide, 1X protease inhibitor). Samples were centrifuged at 14,000 x g for 10 minutes at 4°C and the supernatant was kept for analysis. 10–50% sucrose gradients were prepared on the BioComp instrument. Lysates were loaded onto sucrose gradients and ultra-centrifuged at 4°C at 40,000 rpm for 2 h in an SW-41 Ti rotor (Beckman). After centrifugation, samples were kept on ice and individual polysome profiles were collected using a BioComp Piston Gradient Fractionator with a Triax flow cell. Absorbance at 260 nm was recorded for 3 control and 3 mutant replicates and traces made in GraphPad Prism software.

### Single Cell RNA Sequencing

*<u>Sample preparation and sequencing.</u>* Embryos from a cross of *polr3a^sa907/+^; Tg(fli1a:egfp)* to *polr3a^sa907/+^; Tg(sox10:mRFP)* zebrafish were collected and screened for GFP and RFP at 1 dpf. Fin clips were taken at 2 dpf and genotyped by KASP. Fish were grouped by genotype. At 3 dpf, heads were cut at the pectoral fin. 10 *polr3a^+/+^* heads were pooled for a single wild-type sample and 10 *polr3a^sa907/sa907^* heads were pooled for a single mutant sample. Heads were dissociated in 0.25% trypsin and quenched with stop solution (30% calf serum, 10mM CaCl_2_, in PBS). Dissociated cells were rinsed with dPBS, resuspended in dPBS + 0.4% BSA, and filtered through a 70μm strainer. Single-cell capture, cDNA synthesis, and library preparation were performed by the Genomics Shared Resource at the University of Colorado Anschutz Medical campus. Single-Cell 3’ Gene Expression profile was obtained using the Chromium GEM-X Single-Cell 3’ Library and Gel Bead Kit from 10x Genomics. After cell counting and viability assessment, 40,000 viable flow-sorted cells per sample were loaded on the 10x Genomics Chromium X instrument for single cell capture to capture less abundant cell types in the head. This was followed by amplification of cDNA and library preparation to recover >30,000 cells per sample. An estimated 32,228 cells were captured from the mutant sample with an average of 24,182 reads/cell and an estimated 43,611 cells were captured from the control sample with an average of 16,775 reads/cell. Libraries were quantified, pooled, and sequenced on the Illumina NovaSeq X sequencer at ∼20,000 reads/cell at the University of Colorado - Genomics Shared Resource.

*Data Processing:* Cellranger v7.1.0 (10X Genomics) was used for the reference genome and alignment.

Processing of the sequencing data was performed using Pluto (https://pluto.bio). Feature-barcode matrices generated by 10x Genomics Cell Ranger (10x Genomics Cell Ranger v7.1.0) were obtained for each sample in the dataset. For each sample, transcript count matrices were created by assigning cell barcodes as column names and genes as row names. Transcripts were aggregated to the gene-level by summing counts across duplicated gene names. Individual count matrices from different samples were merged into a single, consolidated matrix. Multiplet class information was added to the cell-level metadata using scDblFinder (v1.14.0), which was applied to each sample independently. Data was processed using the single cell data analysis software Seurat (v4.3.0). An initial Seurat object was created using the CreateSeuratObject() function, where the counts parameter was set to the merged transcript count matrix obtained from the Prepare data step, using a min.cells parameter of 15 and a min.features parameter of 200. To assess the quality of individual cells, the percentage of reads that mapped to mitochondrial genes in each cell was computed. A novelty score was also added to the data, which was derived by taking the logarithm (base 10) of the gene counts divided by the logarithm (base 10) of the UMI counts. Sample-level information, cell-level statistics prepared by Seurat, and multiplet prediction from scDblFinder were incorporated into the Seurat object’s metadata. Cells were removed from the Seurat object based on the following thresholds: UMI counts below 500, UMI counts exceeding 20,000, feature counts below 500, and feature counts exceeding 5,000. Cells with novelty scores below 0.8 or with a percentage of mitochondrial UMIs above 20% were also excluded. Cells identified as potential doublets/multiplets were removed from the data. RNA counts were log-normalized using Seurat’s NormalizeData() function using a scale factor of 10,000 to ensure comparable expression values across cells. Variable features were identified with FindVariableFeatures(), using the variance-stabilizing transformation (VST) method and 3,000 variable features. The cell cycle was scored using the CellCycleScoring() function. The top 3,000 variable features were scaled using the ScaleData() function. Variation due to the difference between G2M and S cell cycle phase scores was regressed out of the top 3,000 variable genes to guide dimensionality reduction and cell clustering. After filtering, 26,392 mutant cells and 34,189 control cells were used for further data processing and analysis. Dimensionality reduction was performed using the top 3,000 variable features on the normalized Seurat object. Principal component analysis (PCA) (RunPCA()) was applied to capture major sources of variability. Uniform manifold approximation and projection (UMAP) (RunUMAP()) was utilized for non-linear dimensionality reduction, using 20 neighboring points and the first 30 principal components. t-distributed stochastic neighbor embedding (t-SNE) (RunTSNE()) was used for further visualization, using the first 30 principal components. To cluster the data, the K-nearest neighbor graph was first computed using the FindNeighbors() function, with k = 20 and the first 30 dimensions of the reduction. Clusters were identified using the FindClusters() function for the following resolution(s): 0.2, 0.6, 1.0, and 1.5.

*<u>Marker gene identification</u>*: Marker genes for each cluster were identified using the RunPrestoAll() function (Presto v1.0.0), with the Wilcoxon rank sum test and a log2 fold change threshold of 0.25. Only positive marker genes for each cluster were returned. For the dot plot, the top 3 marker genes for each cluster were identified by sorting the marker genes for each cluster first by a gene’s adjusted p-value, and then by a gene’s average log2 fold change.

*<u>Marker expression analysis</u>*: Expression data for genes of interest were pulled from the Seurat1 object data slot of the RNA assay, which contains log normalized values. Cell scatter plot showing the expression values for genes across individual cells, using UMAP-defined coordinates. Points are colored by expression. Plots were generated using Pluto (https://pluto.bio).

*<u>Differential expression analysis:</u>* Differential expression analysis was performed with the Seurat R package (R v4.3.1; Seurat v4.3.0), comparing the groups: mutant vs wt and 11 - cartilage - mutant vs 11 - cartilage - wt. Differentially expressed genes were identified using the RunPresto SeuratWrappers function, with the Wilcoxon rank sum test. Genes were only tested if they were detected in a minimum fraction of 0.01 cells in either of the two groups. Log2 fold change was calculated for the comparison mutant vs wt.

*<u>Volcano plot:</u>* Volcano plot showing the log2 fold change of each gene on the x-axis and the - log10(adjusted p-value) on the y-axis. Points are filled if the gene’s adjusted p-value is ≤ 0.01 and its fold change is either less than 0.8 or greater than 1.2. p-value adjustment was performed using Bonferroni correction based on the total number of genes in the dataset. Adjusted p-values are shown on the y-axis of the volcano plot. An adjusted p-value of 0.01 was used as the threshold for statistical significance.

Extremely small adjusted p-values are treated as 0, and for plotting purposes adjusted p-values of 0 are displayed as a maximum -log10 value of 310 on the volcano plot. Analysis was performed and plots generated using Pluto (https://pluto.bio).

*<u>Metascape Analysis</u>*: Differentially expressed genes were exported from Pluto (https://pluto.bio). Genes with an adjusted p-value <0.05 and a fold change greater than 1.2 were included to determine upregulated terms in mutant samples. Genes with an adjusted p-value <0.05 and a fold change less than 0.8 were included to determine downregulated terms in mutant samples. Gene lists were entered into Metascape [60] for pathway analysis.

### Statistics

All statistical analysis was performed using GraphPad Prism software (version 10.4.0). For comparisons between two groups (control vs mutant), a two tailed t-test with Welch’s correction was performed to account for unequal sample numbers. For comparisons across multiple groups, the Brown-Forsythe and Welch ANOVA tests were performed followed by Dunnett’s T3 multiple comparisons test to determine statistical significance between groups. All graphs show the mean ± standard deviation. For all comparisons, p-values were calculated in GraphPad Prism and p<0.05 was determined statistically significant.

## Acknowledgements

The authors thank Dr. Paul Trainor for comments on the research; Christine Archer, Olivia Gomez, and Paige Mashman for zebrafish care and maintenance; and Emma Heiny for genotyping assistance. We would like to thank the personnel from the University of Colorado Anschutz Medical Campus Advanced Light Microscopy Core for imaging assistance. Imaging was performed in the Advanced Light Microscopy Core facility of the NeuroTechnology Center at the University of Colorado Anschutz Medical Campus, which is supported in part by Rocky Mountain Neurological Disorders Core Grant (P30NS048154) and by Diabetes Research Center Grant (P30 DK116073). We also thank the personnel from the Genomics Shared Resource for single cell capture, library preparation, and sequencing. The Genomics Shared Resource at the University of Colorado Anschutz Medical Campus is supported by the Cancer Center Support Grant (P30CA046934).

## Supporting Information

**S1 Fig. *polr3a^sa907^* is in a highly conserved region of POLR3A and leads to a reduction in Polr3a expression.** (A) The POLR3A/Polr3a protein sequence is conserved across humans, zebrafish, and other vertebrates at the site of the sa907 mutation (yellow). Locations of variants associated with POLR3-HLD (grey) and WRS (light blue) are also indicated. Amino acids 541-660 are represented here. Full length POLR3A has 1390 amino acids. (B) Sanger sequencing shows the sa907 allele is a C>T nonsense mutation in exon 14. (C) Western blotting shows a reduction in full length Polr3a in *polr3a^sa907/sa907^* zebrafish at 5 dpf (155 kDa, black arrowhead). Mutants may produce a truncated protein (78 kDa, magenta arrowhead). α-tubulin was used as a loading control. Quantification shows at least a 90% reduction in Polr3a protein in *polr3a^sa907/sa907^*zebrafish.

**S2 Fig. Craniofacial cartilage and bone are hypoplastic in *polr3a^sa907/sa907^* zebrafish**. (A-F) Alizarin red and Alcian blue staining at 6 dpf reveals differences in the size of skeletal elements in mutant zebrafish. Ventral views (B,E) show overall head length (G, ***p = 0.0002) and eye diameter (H, ****p <0.0001) are smaller (measurements indicated in panel B). The length of the ceratohyal (ch) is shorter in mutants (I, *** p = 0.0005), as is the width of Meckel’s (J, **p = 0.003) from the left to right jaw joint (measurements indicated in panel E). Abbreviations: M, Meckel’s cartilage; ch, ceratohyal. Scale bar = 100 µm. n = 8 controls, n = 8 mutants.

**S3 Fig. Craniofacial cartilage and bone remain hypoplastic in *polr3a^sa907/sa907^* zebrafish at 8 dpf**. (A-H) Alizarin red and Alcian blue staining at 8 dpf demonstrates the overall reduction of skeletal elements in mutant zebrafish. (A,B) Lateral views show the size difference between control and mutant larvae. (C,D) Flat mounts of the neurocranium show reductions in cartilage formation, especially around the eye. (E,F) Flat mounts of the viscerocranium reveal the reduction of NCC-derived cartilage and bone structures including the maxilla (mx), ceratohyal (ch), and Meckel’s cartilage (M). (G,H) Dissection of arch 1 and 2-derived elements show the differences in bone at 8 dpf. The opercle (op) remains reduced in size, there is a small amount of bone forming in the entopterygoid (en) in mutants, while other bones fail to form. Overall head length (I, ***p <0.0001) and eye diameter (J, ***p = 0.0003) are smaller. The length of the ceratohyal (ch) is shorter in mutants (I, *** p = 0.0004), as is the width of Meckel’s (J, *p = 0.048) from the left to right jaw joint (as in S2 Fig). Scale bar = 200 µm. n = 4 controls, n = 4 mutants.

**S4 Fig. Pharyngeal arches develop normally in *polr3a^sa907/sa907^* embryos.** (A,B) *sox10:egfp* expression in control and *polr3a^sa907/sa907^* zebrafish at 36 hpf. Pharyngeal arch 1 and 2 are outlined and quantification of this region shows no significant difference between controls and mutants (C). (D,E) *sox10:egfp* expression at 48 hpf also shows no significant changes in arch volume, outlined area (F). Scale bar = 100 µm.

**S5 Fig. In situ hybridization for *dlx2a* does not show significant differences between control and *polr3a^sa907/sa907^* embryos.** At 48 hpf, no differences were detected in *dlx2a* expression, as a marker of the pharyngeal arches. White arrowheads indicate anatomical location of arch 1 and 2. Scale bar = 100 µm.

**S6 Fig. Quantification of cell morphology in Meckel’s cartilage does not reveal significant differences in *polr3a^sa907/sa907^* embryos at 5 dpf.** (A,B) Images of Meckel’s cartilage using *sox9a:egfp* expression. (C,D) Cells in Meckel’s cartilage were outlined using CellProfiler. (E,F) Representation of cell masking from CellProfiler showing the organization of cells. These masks were used to quantify cell number, area, and eccentricity. There were no differences in the average area per cell (G; p = 0.478) or in cell eccentricity (H; p = 0.196) in *polr3a^sa907/sa907^* mutants compared to controls. n = 18 controls, n = 13 mutants.

**S7 Fig. In situ hybridization for *col2a1a* reveals that early cartilage differentiation is unaffected in *polr3a^sa907/sa907^* zebrafish at 2 dpf and 3 dpf.** Staining in the region of arch 1 and 2-derived cartilage is unaffected (arrows). Scale bar = 100 µm.

**S8 Fig. Early bone development is unchanged in *polr3a^sa907/sa907^* zebrafish.** (A,B) *sp7:gfp* to label osteoblasts (green) and Alizarin red for bone (magenta) shows no significant changes at 3 dpf in either *sp7:gfp* (C) or Alizarin red (D). n = 19 controls, n = 10 *polr3a^sa907/sa907^*. Scale bar = 100 µm.

**S9 Fig. Bone development remains reduced though 8 dpf in *polr3a^sa907/sa907^* zebrafish.** A, B) *sp7:gfp* to label osteoblasts (green) and Alizarin red for bone (magenta) shows a significant reduction in Alizarin red volume in the opercle (op) in mutant larvae (C; ****p < 0.0001). n = 39 controls, n = 11 *polr3a^sa907/sa907^*. (D,E) Ventral views demonstrate persistence of bone hypoplasia in mutant zebrafish in all craniofacial bones examined. Quantification of the dentary (de; F; n = 15 controls, n = 7 mutants), maxilla (mx, G; n = 15 controls, n = 9 mutants), and branchiostegal rays (bsr, H; n = 13 controls, n = 8 mutants) further demonstrate that these elements remain significantly reduced. ****p < 0.0001. The maxilla was not detected in 3 larvae examined (noted by the * in D). The entopterygoid (en) remains smaller in mutant larvae. By 8 dpf, we detect little endochondral ossification in the ceratohyal (ch) in mutants, noted by the + in panel D. Scale bar = 100 µm.

**S10 Fig. *polr3a^sa907/sa907^* zebrafish do not show differences in Ca^2+^ regulation.** (A-C) Expression of corpuscle marker *cdh16* is unchanged in *polr3a* mutants relative to controls. ns, not significant. (D-F) *polr3a^sa907/sa907^* mutant zebrafish are hyposensitive to acoustic stimuli. ****p < 0.0001; *p = 0.024. n = 37 controls, n = 25 *polr3a^sa907/sa907^*. Scale bar = 50 µm.

**S11 Fig. *polr3a* expression during development.** (A) qRT-PCR expression of *polr3a* in wild type embryos shows extensive maternal expression at 4-cell stage, and then increasing expression from 3-5 dpf. (B-D) HCR in situ for *polr3a* shows a broad low-level of expression at 24 hpf (B), 48 hpf (C), and 72 hpf (D), in wild-type embryos. The autofluorescence negative controls do not show expression at any stage (E-G). Scale bar = 100 µm

**S12 Fig. *mbp:tagRFP* expression at 6 dpf is unchanged in *polr3a^sa907/sa907^* zebrafish.** (A,B) *mbp:tagRFP* expression (magenta) is unchanged in the hindbrain (n = 12 controls; n = 12 mutants). (C,D) No changes in *mbp:tagRFP* expression in spinal cord were observed (n = 8 controls; n = 10 mutants) in *polr3a^sa907/sa907^* zebrafish. Quantification of *mbp:tagRFP* volume in the head (E) and spinal cord (F) shows no statistically significant differences. ns, not significant. Scale bar = 100 µm.

**S13 Fig. Analysis of TUNEL and pH3 at 2 dpf within the pharyngeal arches.** (A,B) TUNEL (magenta) and pH3 (white) within the region of arch 1 and 2 (white outline) show no significant differences between control and *polr3a^sa907/sa907^* embryos. (C) Quantification of TUNEL spots within arches 1 and 2 shows no significant difference, nor does pH3 (D). ns, not significant. n = 12 controls, n = 11 *polr3a^sa907/sa907^*. Scale bar = 100 µm.

**S14 Fig. TUNEL and pH3 are unchanged at 2 dpf.** (A,B) Analysis of TUNEL (magenta) and pH3 (white) throughout the head (DAPI, all nuclei; blue outline) at 2 dpf. (C) Quantification of TUNEL and pH3 (D) reveals no significant differences between control and *polr3a^sa907/sa907^* embryos. n = 12 controls, n = 11 *polr3a^sa907/sa907^*. Scale bar = 100 µm.

**S15 Fig. qRT-PCR at 2 dpf shows diminished pre-tRNA transcription.** qRT-PCRs for pre-tRNAs, 7SL RNA, and 5’ETS rRNA shows that only pre-tRNAs are significantly reduced at 2 dpf. ****p <0.0001; ***p = 0.0002.

**S16 Fig. Cluster identification**. (A) Dot plot of top three marker genes for each cluster. (B) UMAP plot of *elavl3* expression as a marker of neuronal cells. (C) UMAP plot of *col2a1a* expression as a marker of cartilage cells. (D) UMAP plot of *sox10* expression. (E) UMAP plot of *sox9a* expression.

**S17 Fig. Metascape analysis of differentially expressed genes between controls and *polr3a^sa907/sa907^* zebrafish.** Downregulated terms include ribosome large subunit biogenesis and cell cycle (blue boxes). Terms upregulated in mutant zebrafish include Tp53 signaling (magenta box).

**S18 Fig. UMAP plots of gene expression for genes in the integrated stress response pathway.** The *polr3a^+/+^* sample (wt) is on the left and the *polr3a^sa907/sa907^* sample (mutant) is on the right. (A) Expression of *atf4b*. (B) Expression of *ddit3.* (C) Expression of *chac1.* (D) Expression of *aars1*.

**S19 Fig. UMAP plots of gene expression for genes in the Tp53 pathway.** (A) UMAP plots of *tp53* expression and (B) *mdm2* expression in wild-type (wt, left) versus *polr3a^sa907/sa907^* zebrafish (right).

**S20 Fig. Proliferation is unchanged in *polr3a^sa907/sa907^* zebrafish at 3 dpf regardless of *tp53* status.**

**S21 Fig. Alcian blue and alizarin red staining at 6 dpf reveals subtle improvements in *polr3a^sa907/sa907^* zebrafish upon *tp53* inhibition.** (A,B) Control zebrafish show bone staining in the opercle and organized stacking in Meckel’s cartilage. *polr3a^sa907/sa907^; tp53^+/+^* zebrafish have smaller opercles (C,D) as do *polr3a^sa907/sa907^; tp53^+/-^* (E,F) and *polr3a^sa907/sa907^; tp53^-/-^* (G,H) zebrafish. Quantification of the width of Meckel’s cartilage (vertical yellow line in B) reveals no significant changes (I). Quantification of ceratohyal length (J), demonstrates no significant improvements in length regardless of *tp53* status. Adjusted p-values: ns, not significant. *p = 0.04; ****p < 0.0001. (K) Head length remains reduced in *polr3a^sa907/sa907^* zebrafish with inhibition of *tp53*. Adjusted p-values: ns, not significant; *p = 0.01; **p = 0.004. n = 24 control, n = 4 *pol3a^sa907/sa907^; tp53^+/+^;* n = 9 *polr3a^sa907/sa907^; tp53^+/-^*; n = 7 *polr3a^sa907/sa907^; tp53^-/-^*. ch, ceratohyal; M, Meckel’s cartilage; op, opercle. Scale bar = 100 µm.

**S22 Fig. Alcian blue and Alizarin red staining at 8 dpf reveals subtle improvements in *polr3a^sa907/sa907^* zebrafish upon *tp53* inhibition.**

