## Supplementary figures and images for "RNA Polymerase III subunit Polr3a is required for craniofacial cartilage and bone development in zebrafish"

### S1 Fig

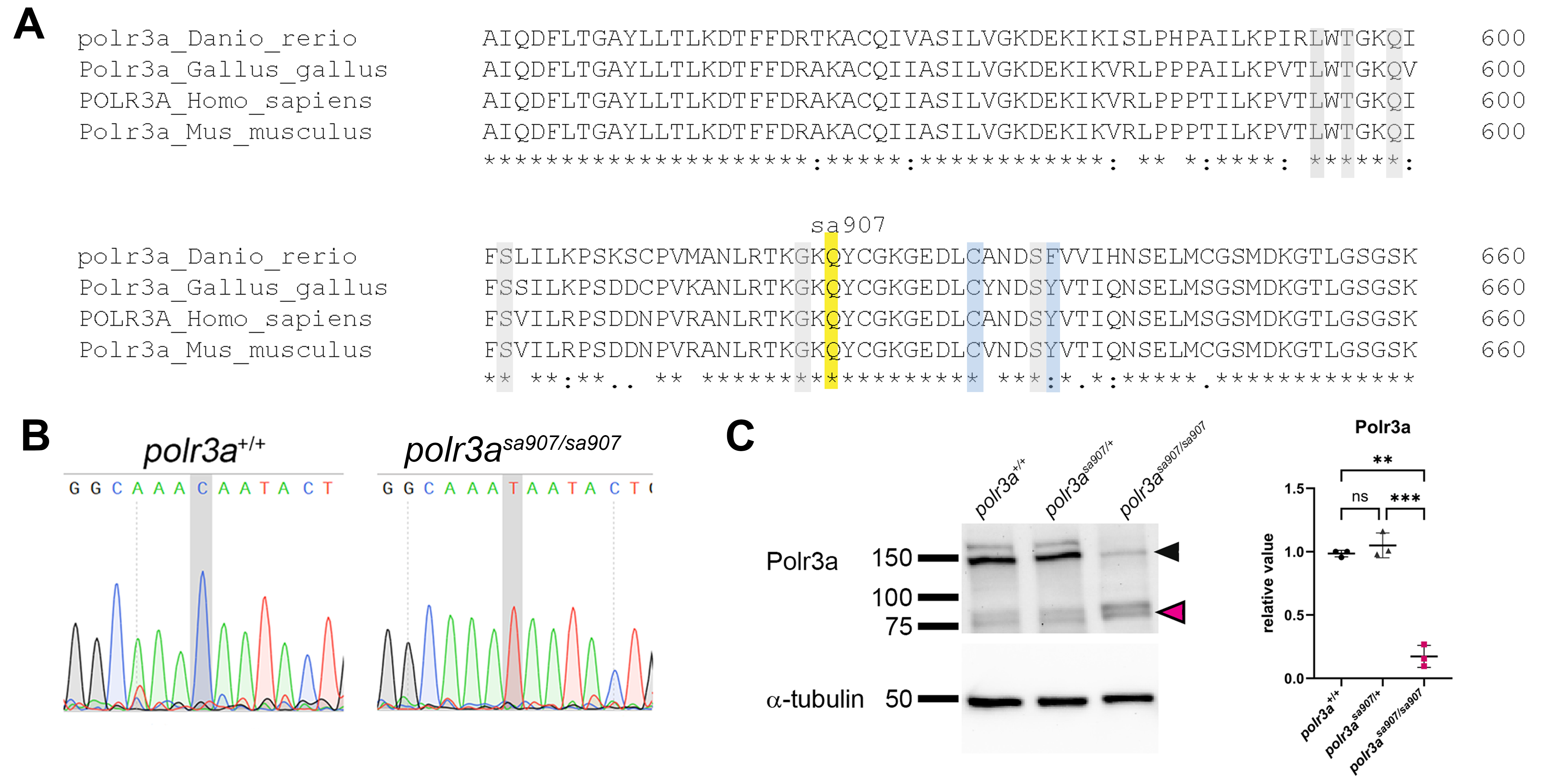

### S2 Fig

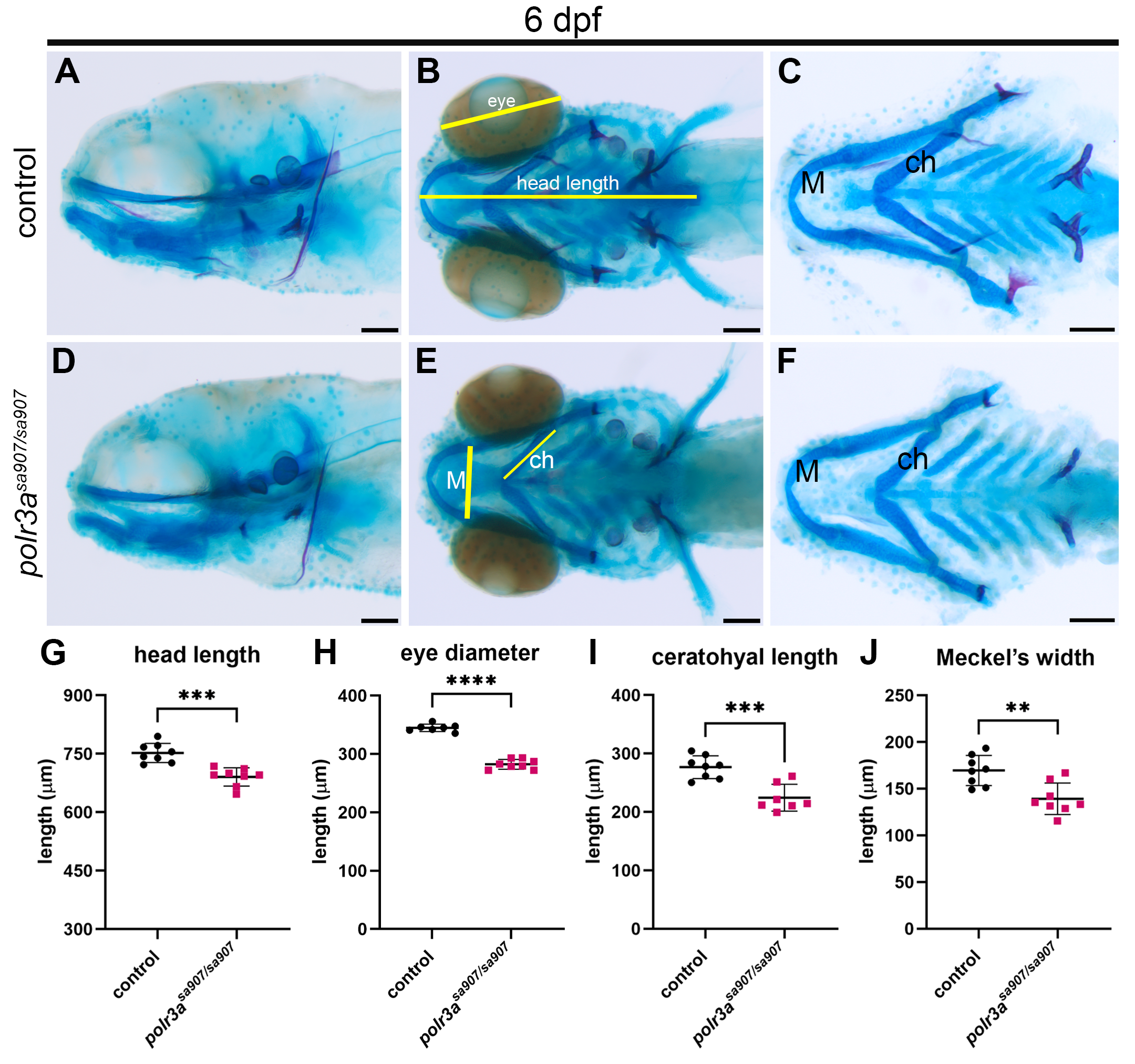

### S3 Fig

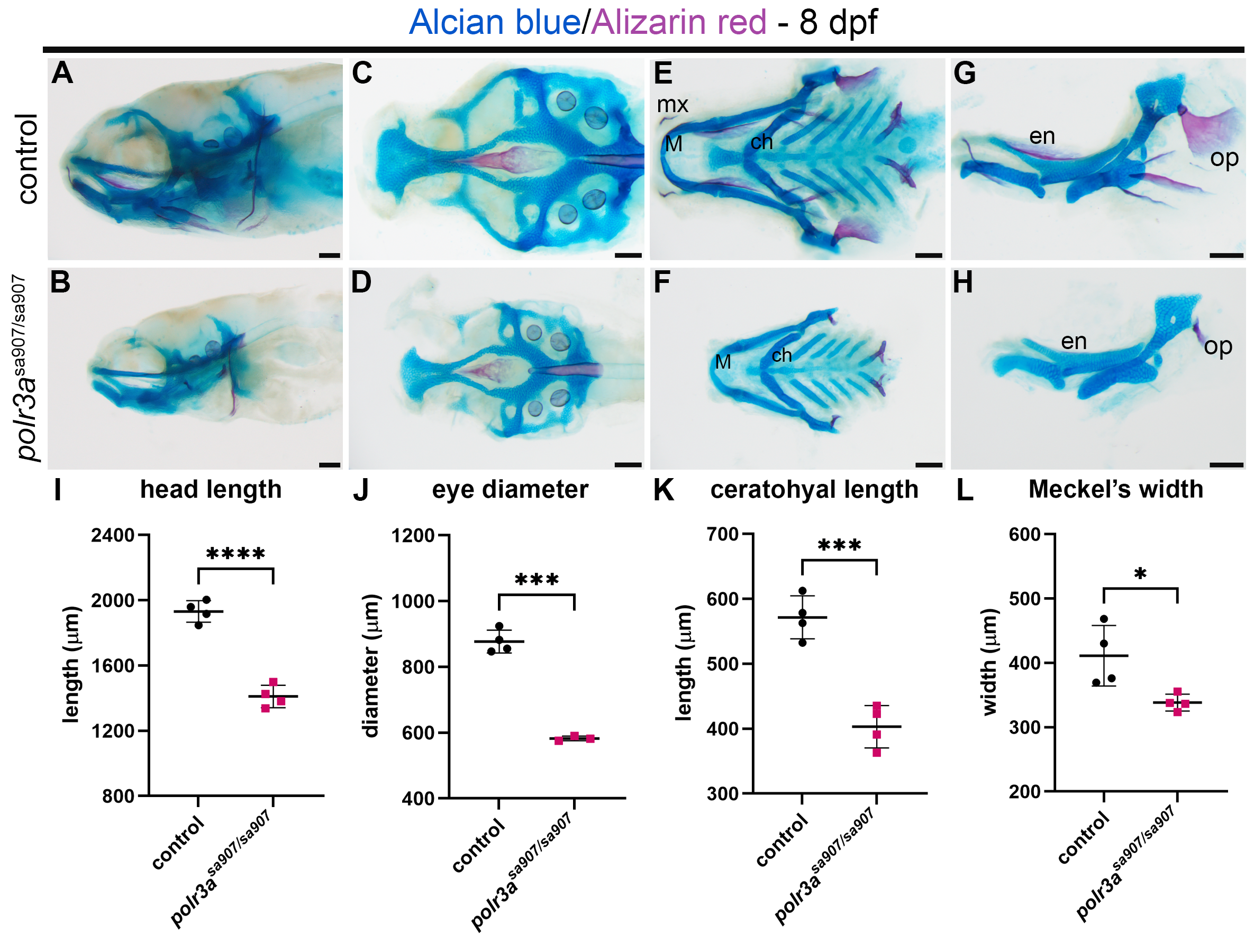

### S4 Fig

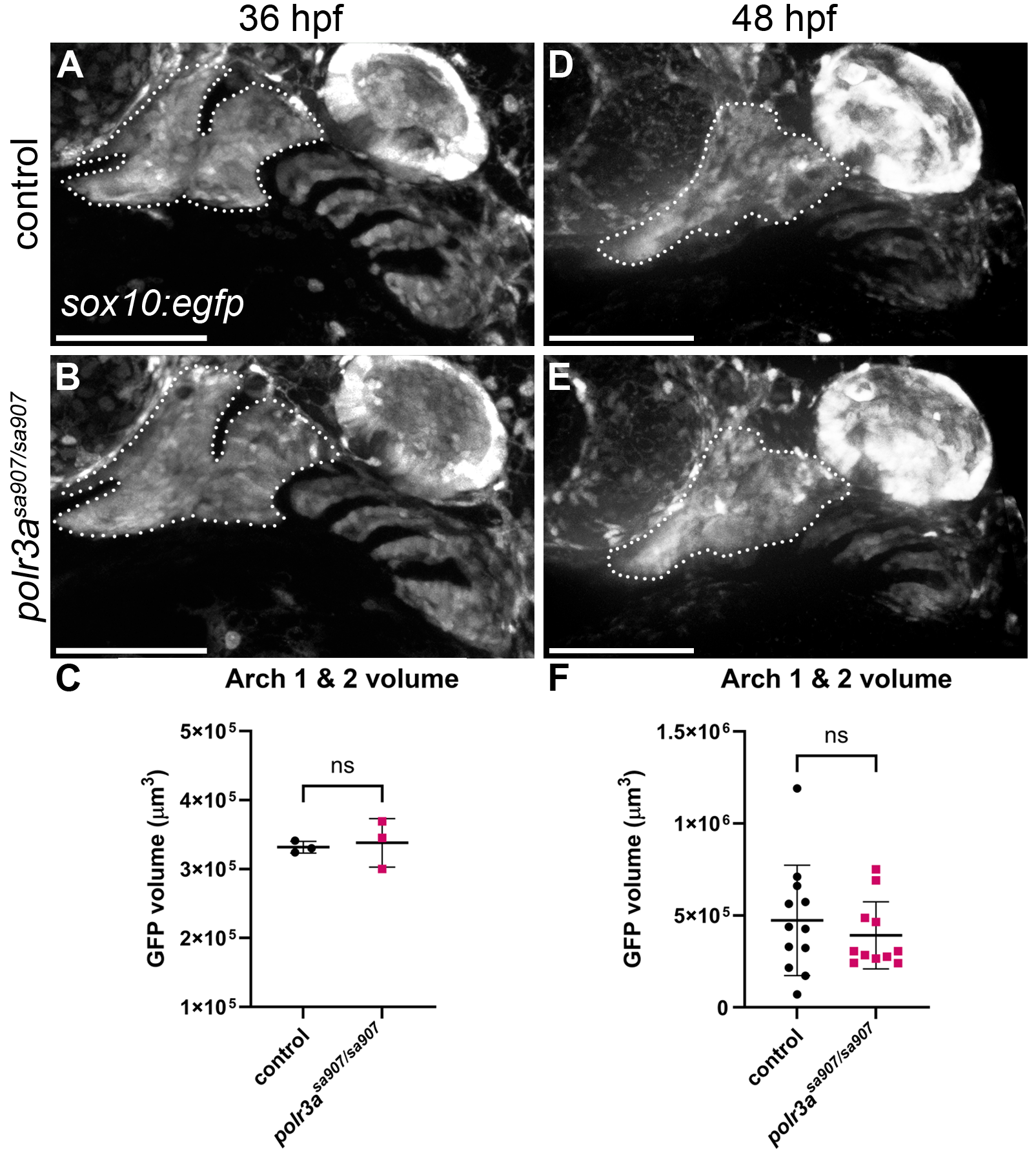

### S5 Fig

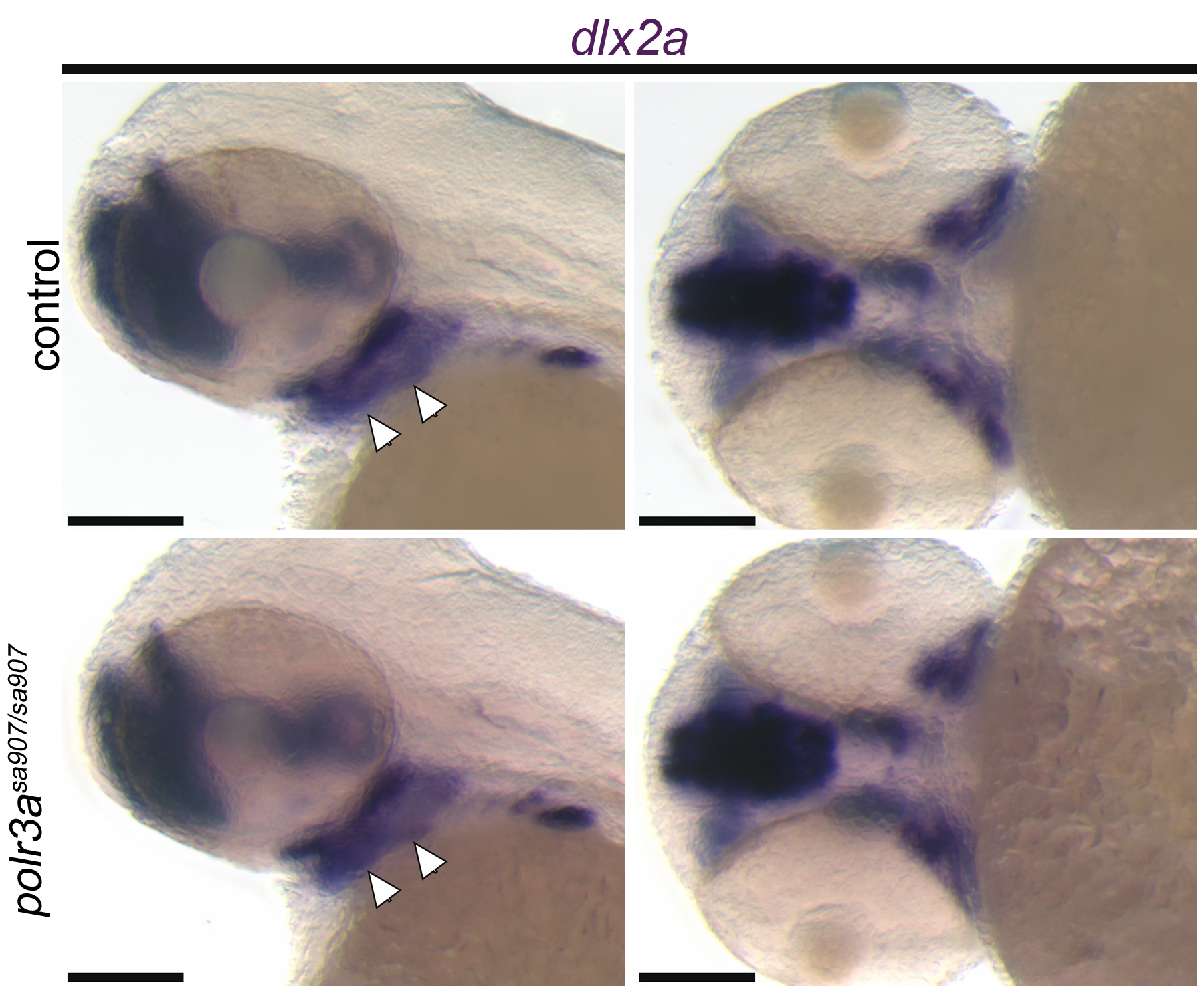

### S6 Fig

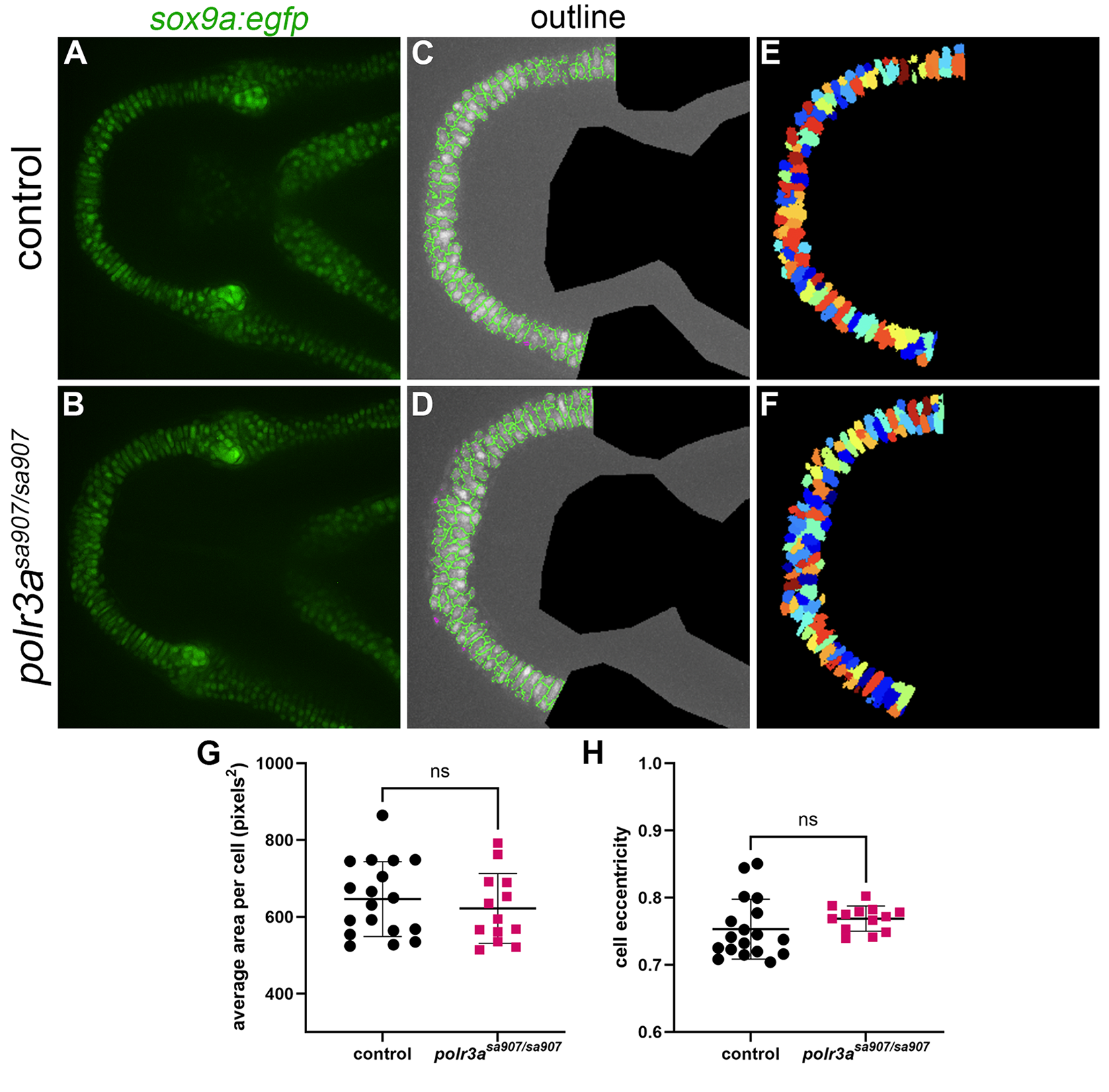

### S7 Fig

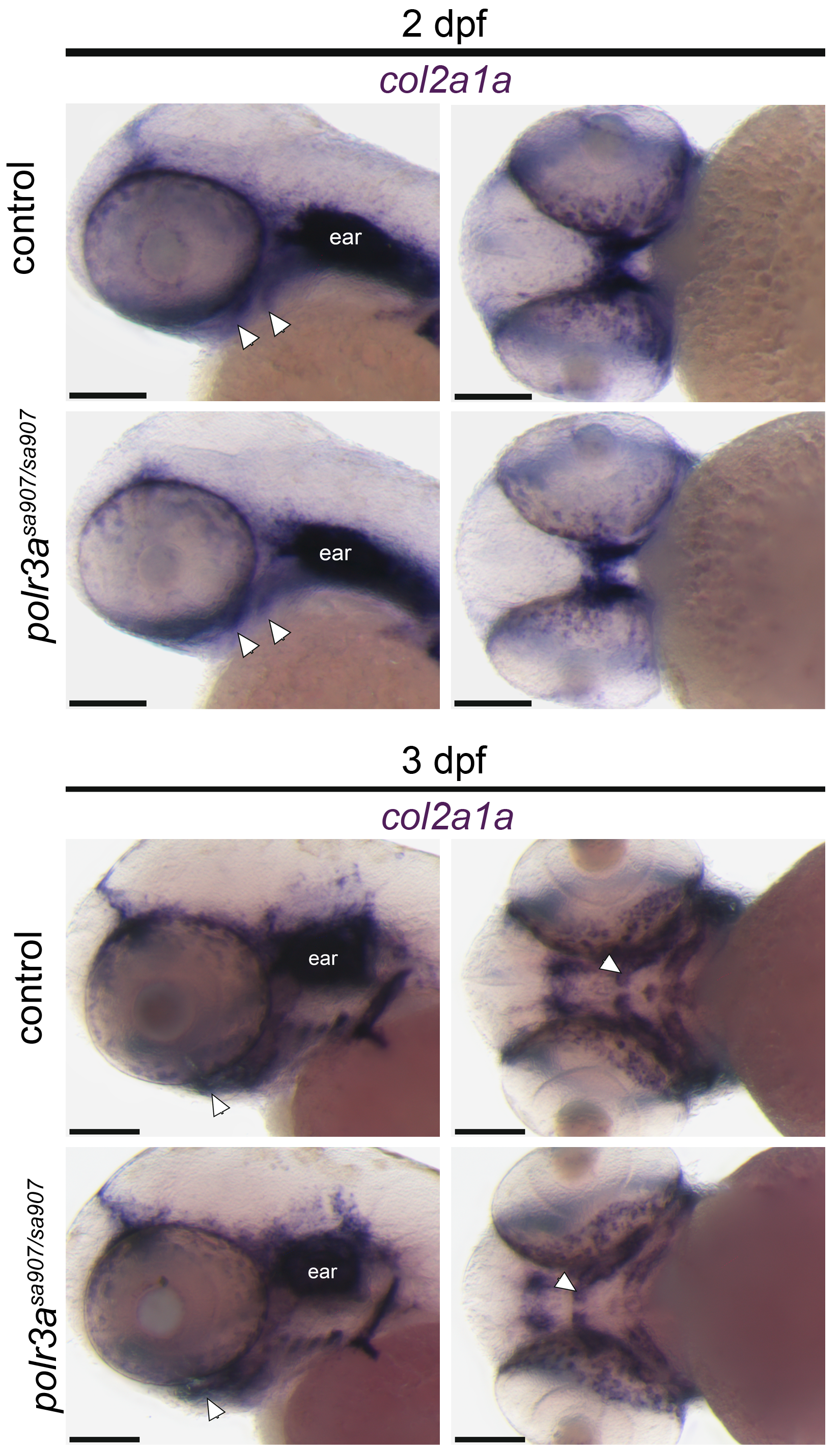

### S8 Fig

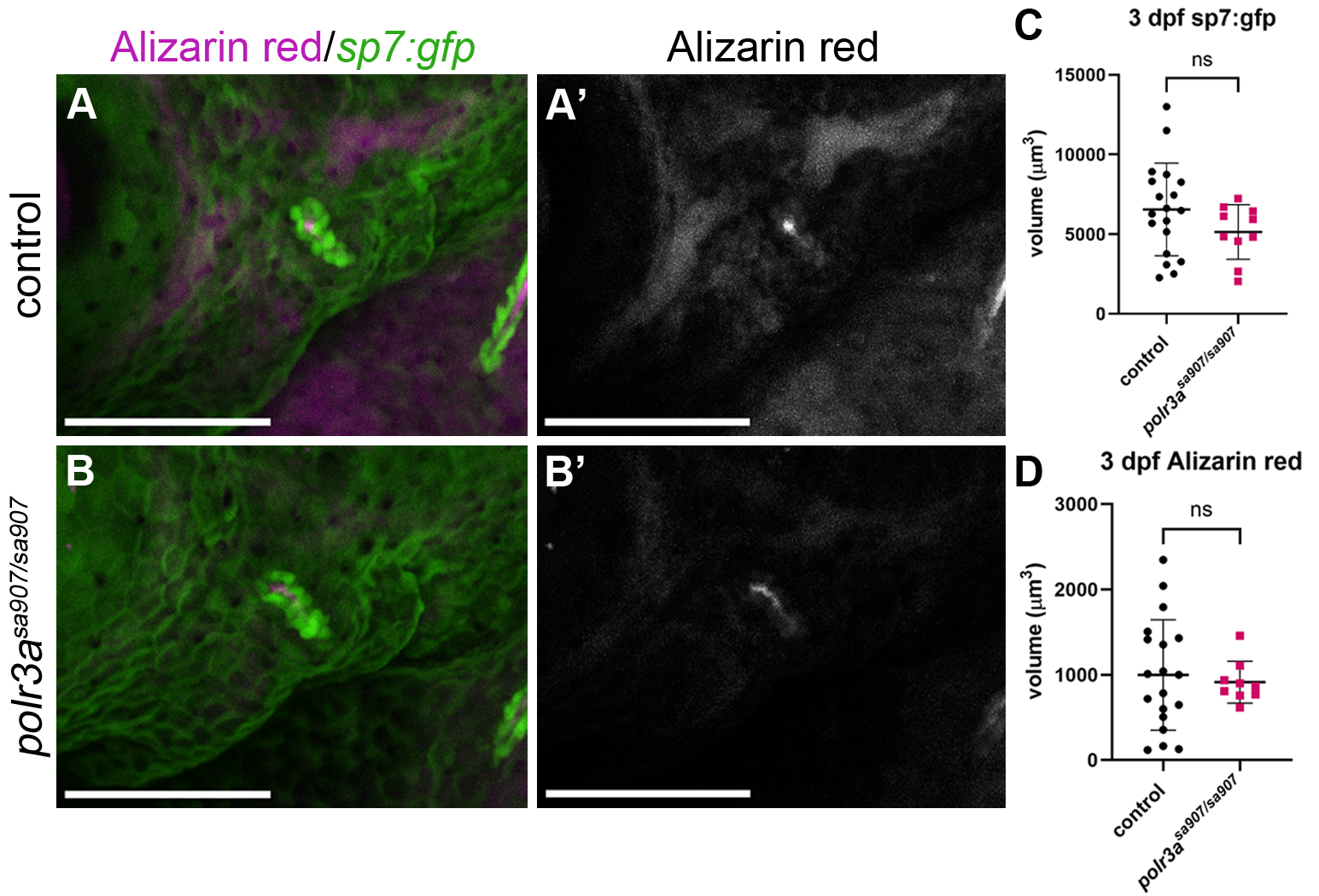

### S9 Fig

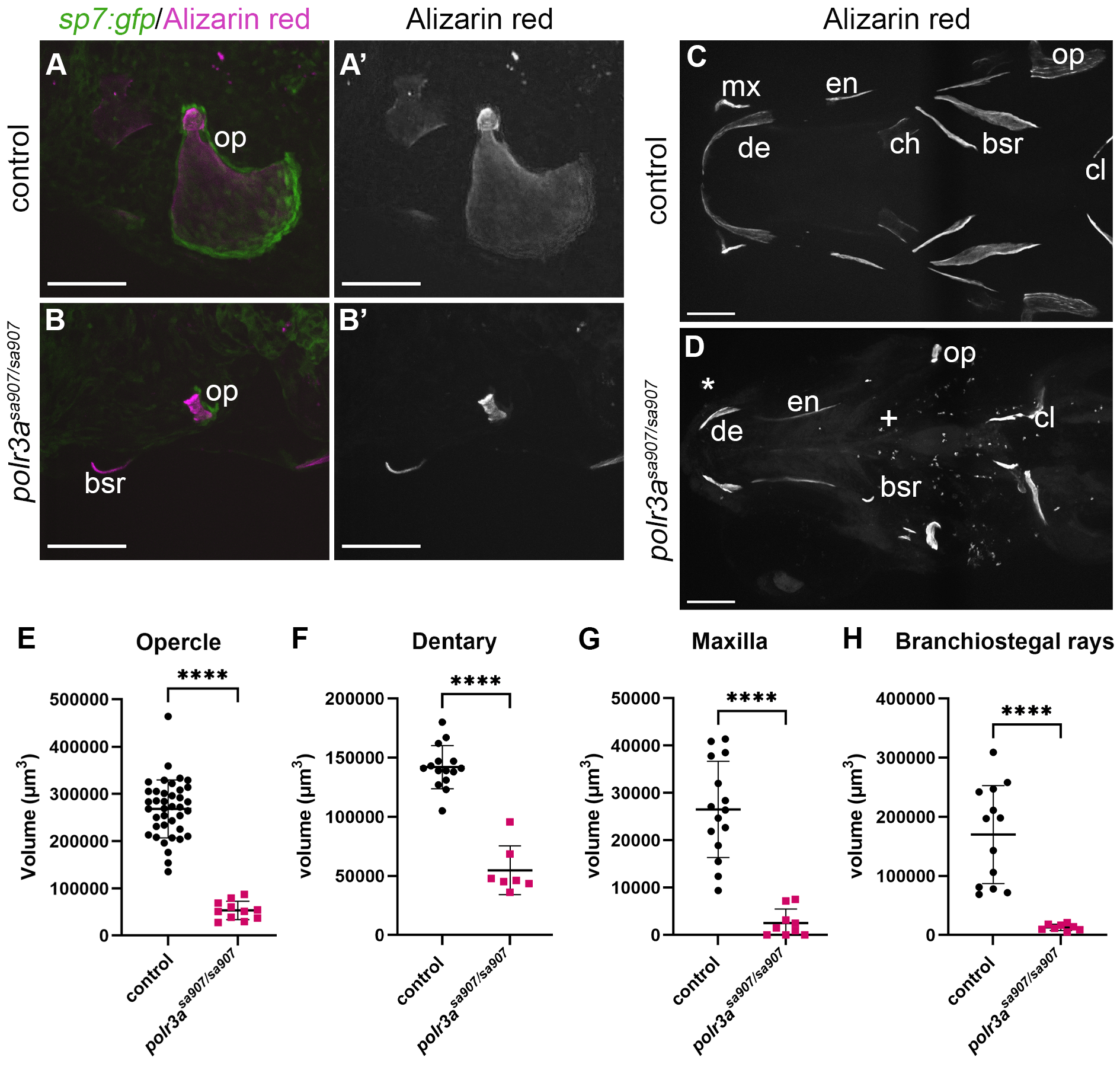

### S10 Fig

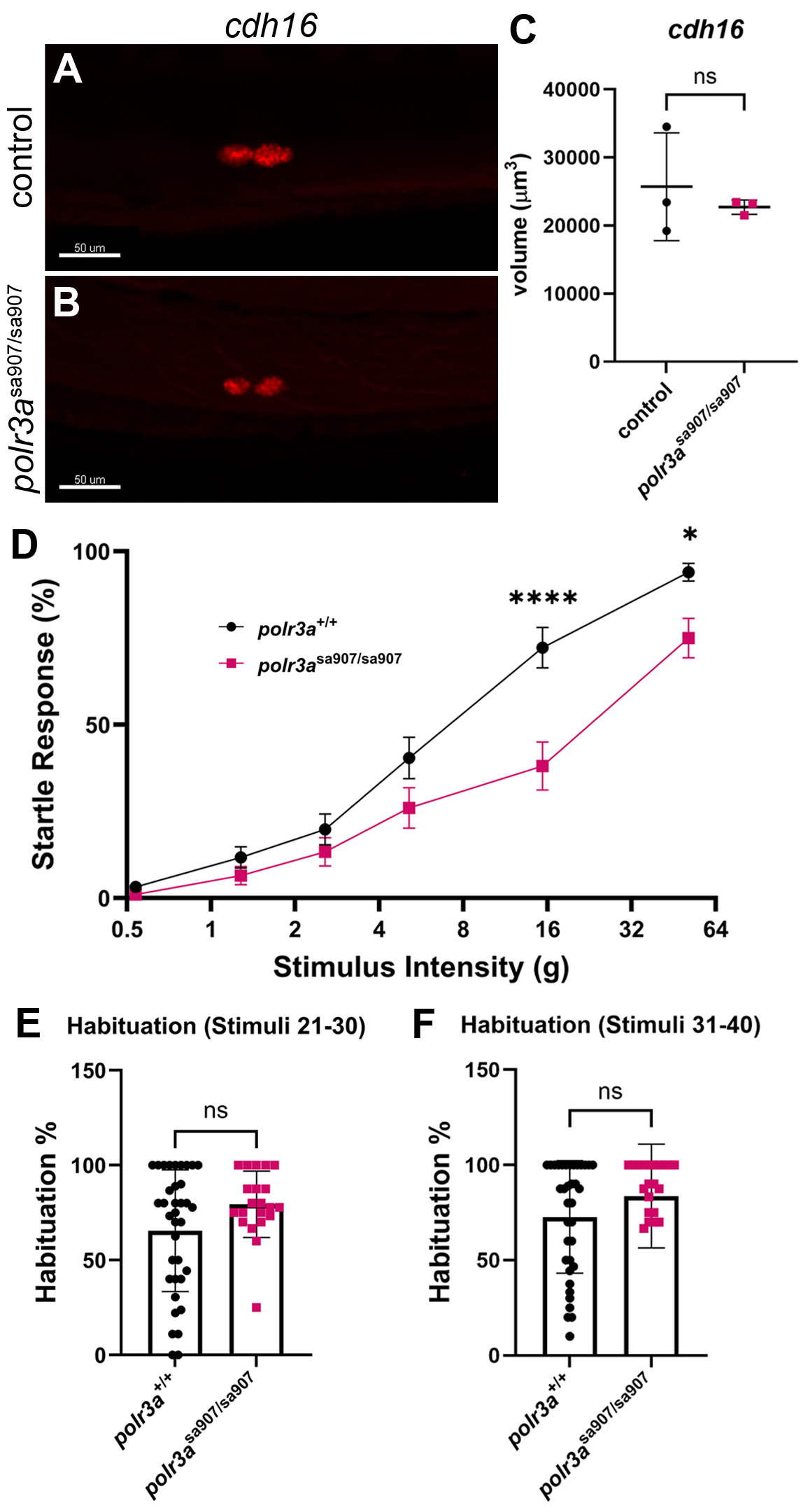

### S11 Fig

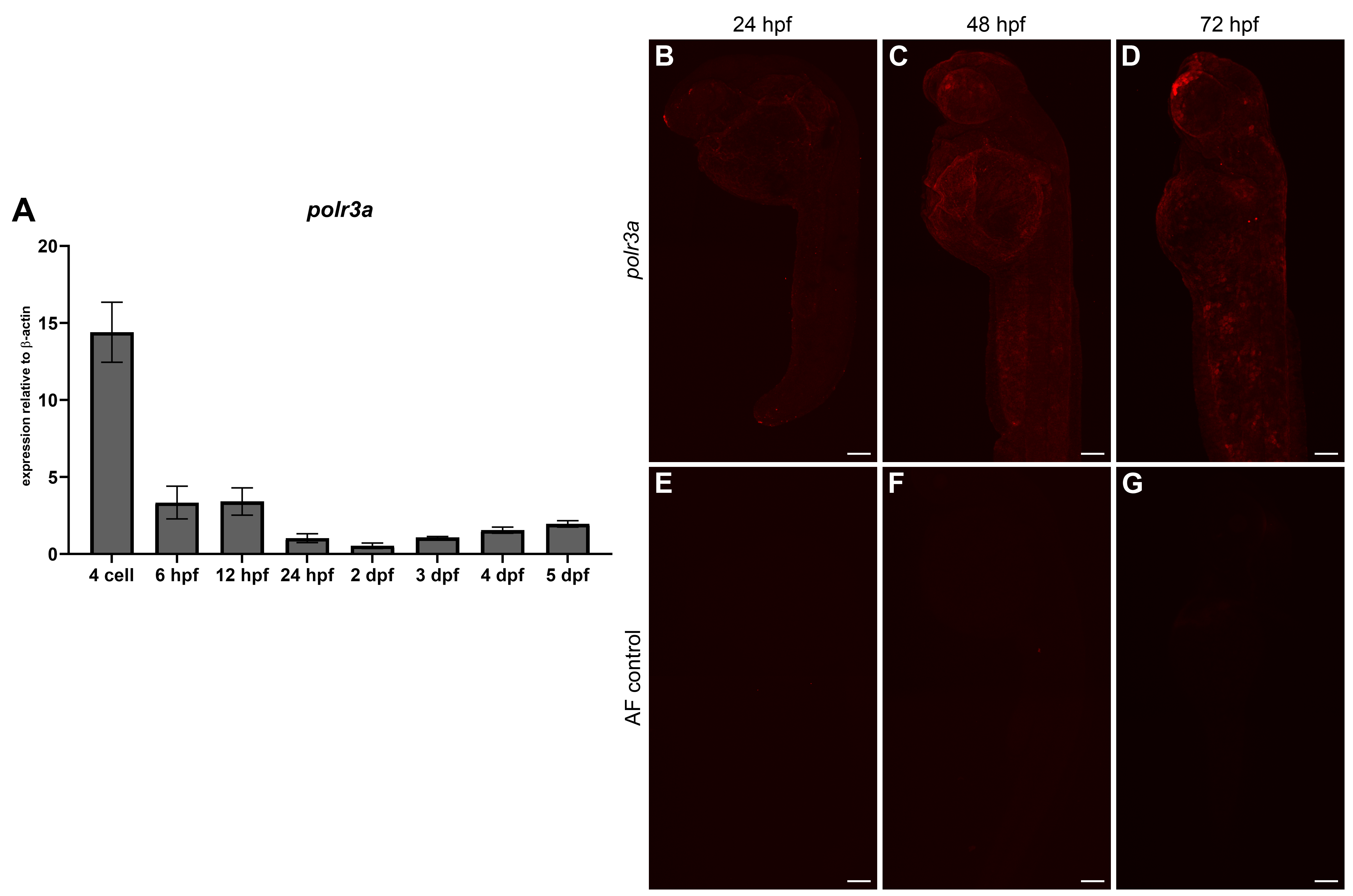

### S12 Fig

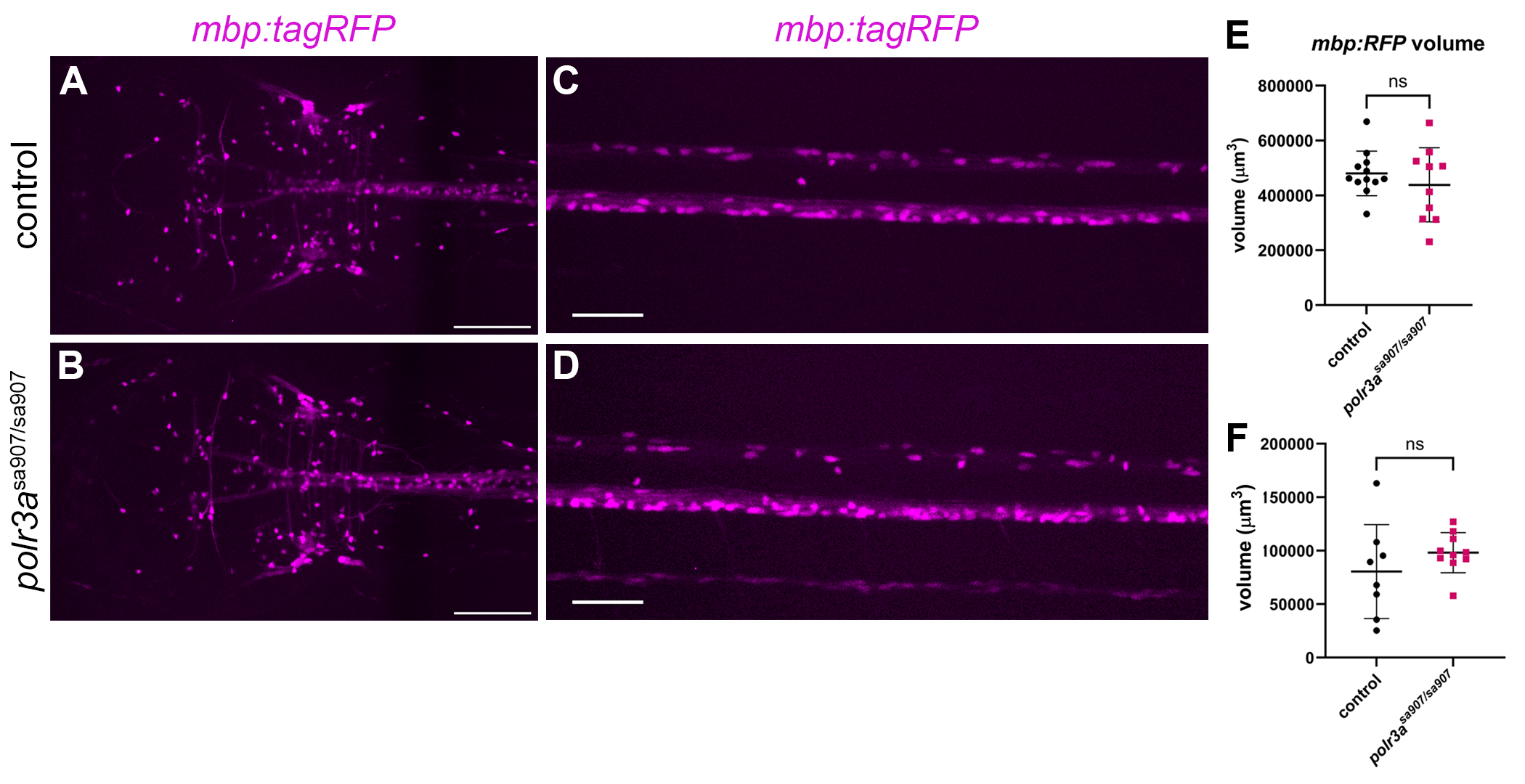

### S13 Fig

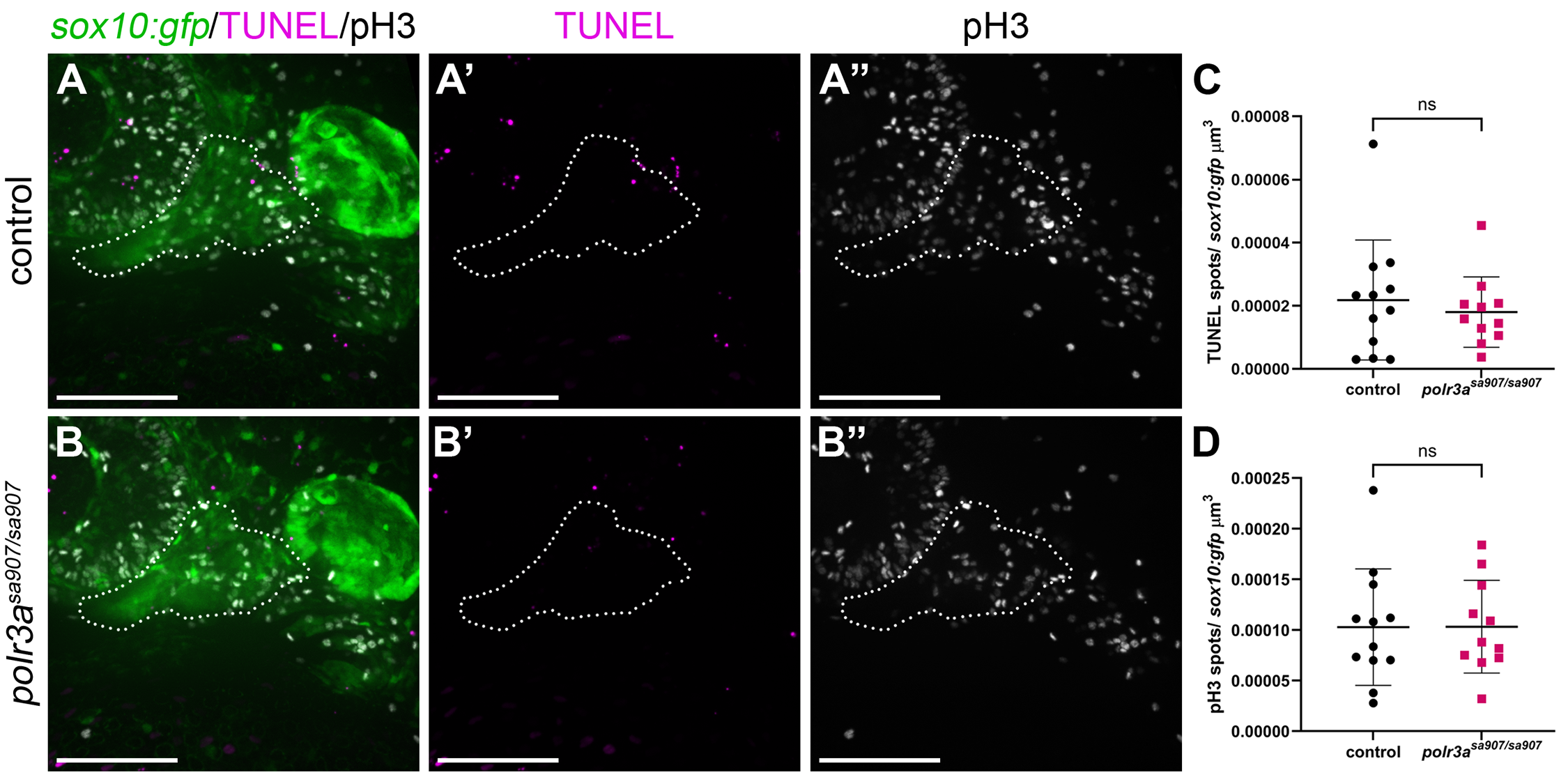

### S14 Fig

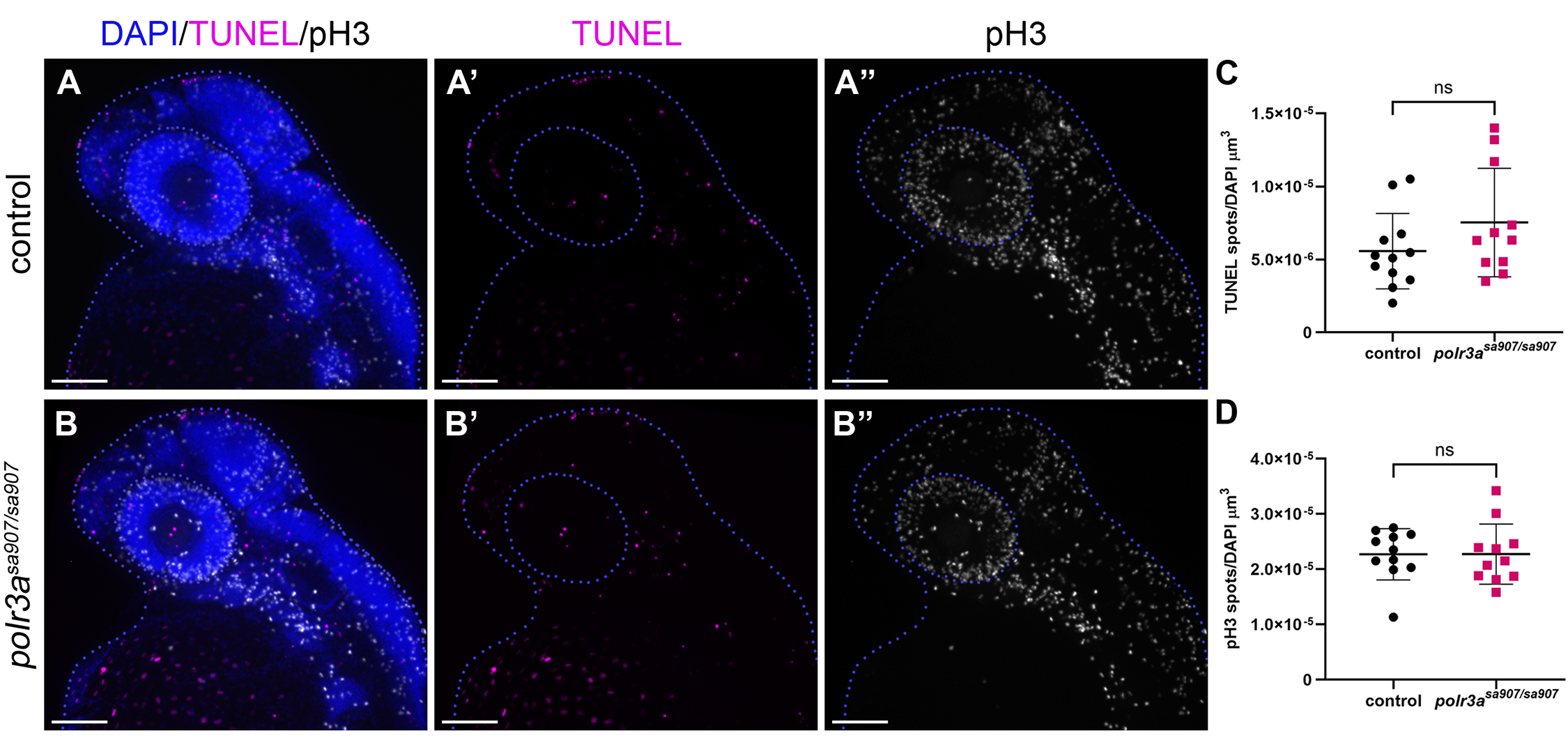

### S15 Fig

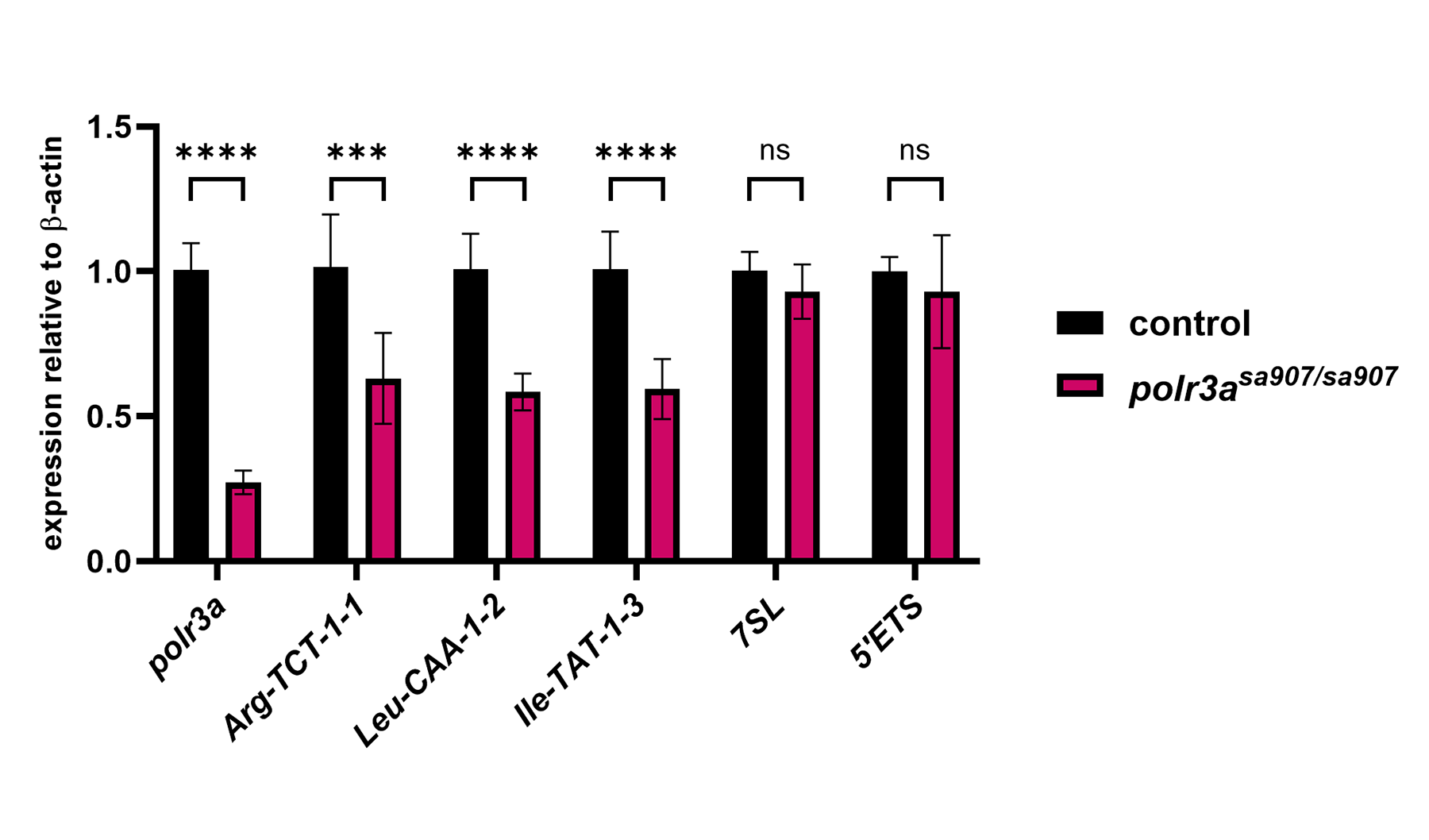

### S16 Fig

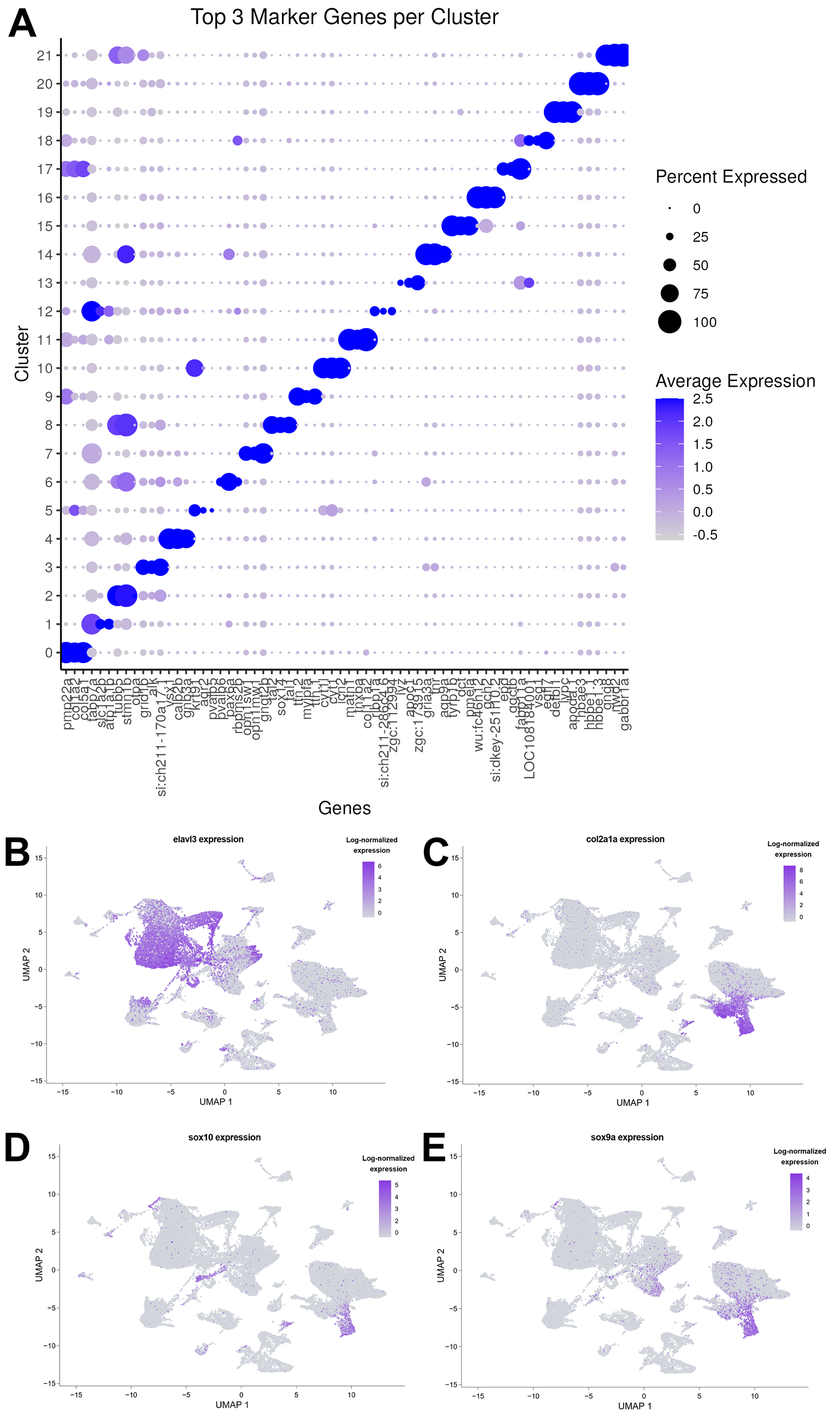

### S17 Fig

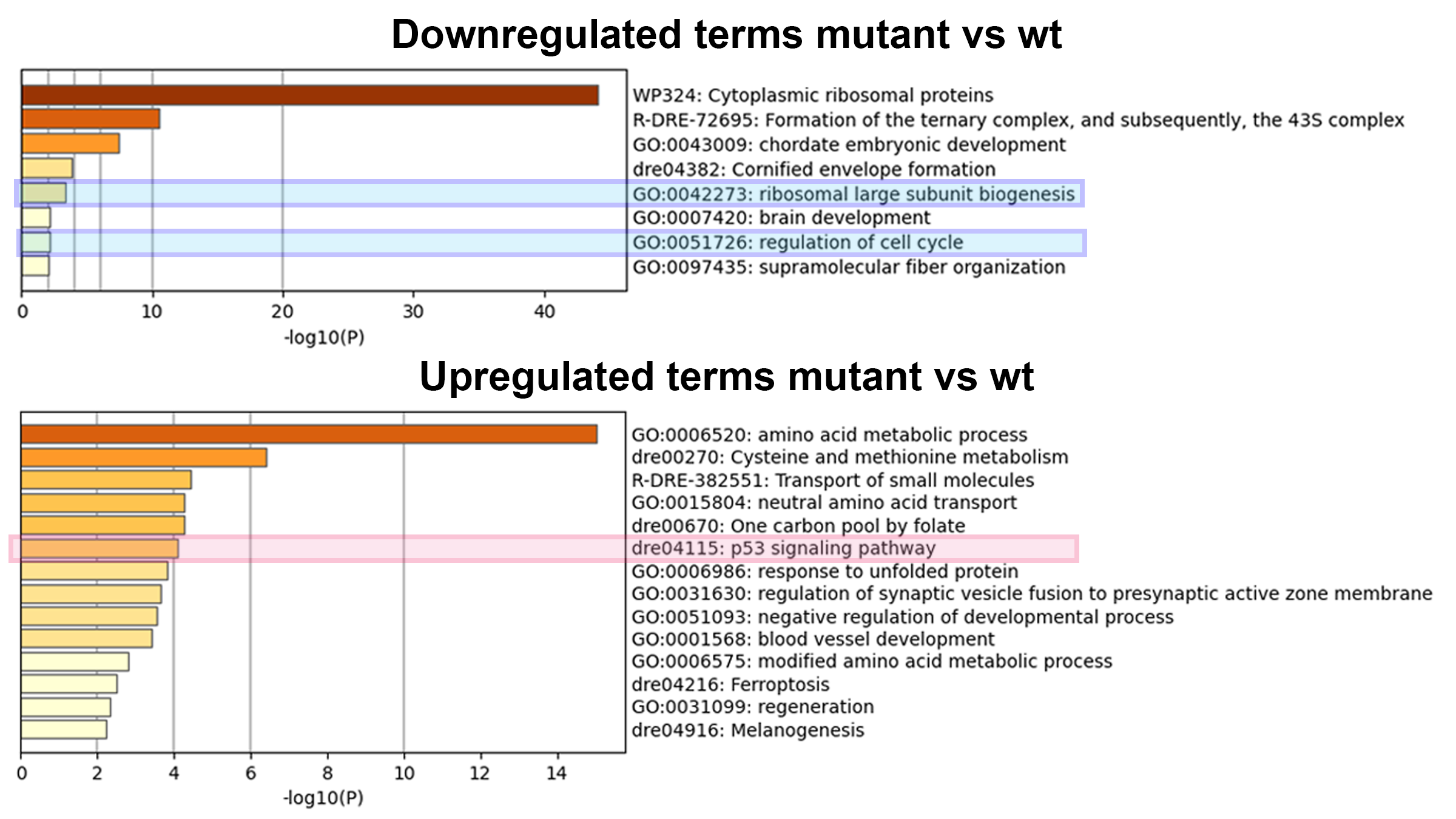

### S18 Fig

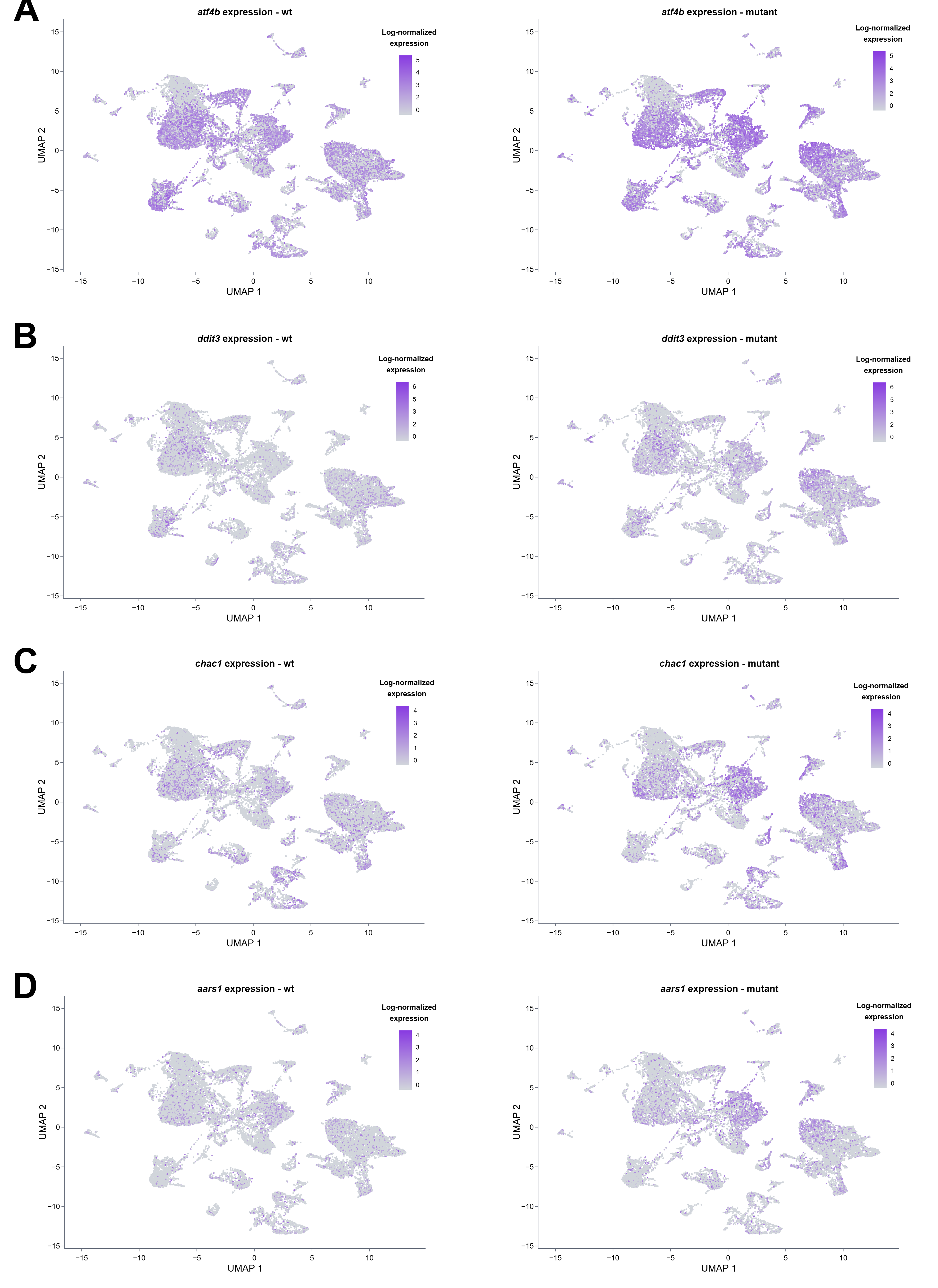

### S19 Fig

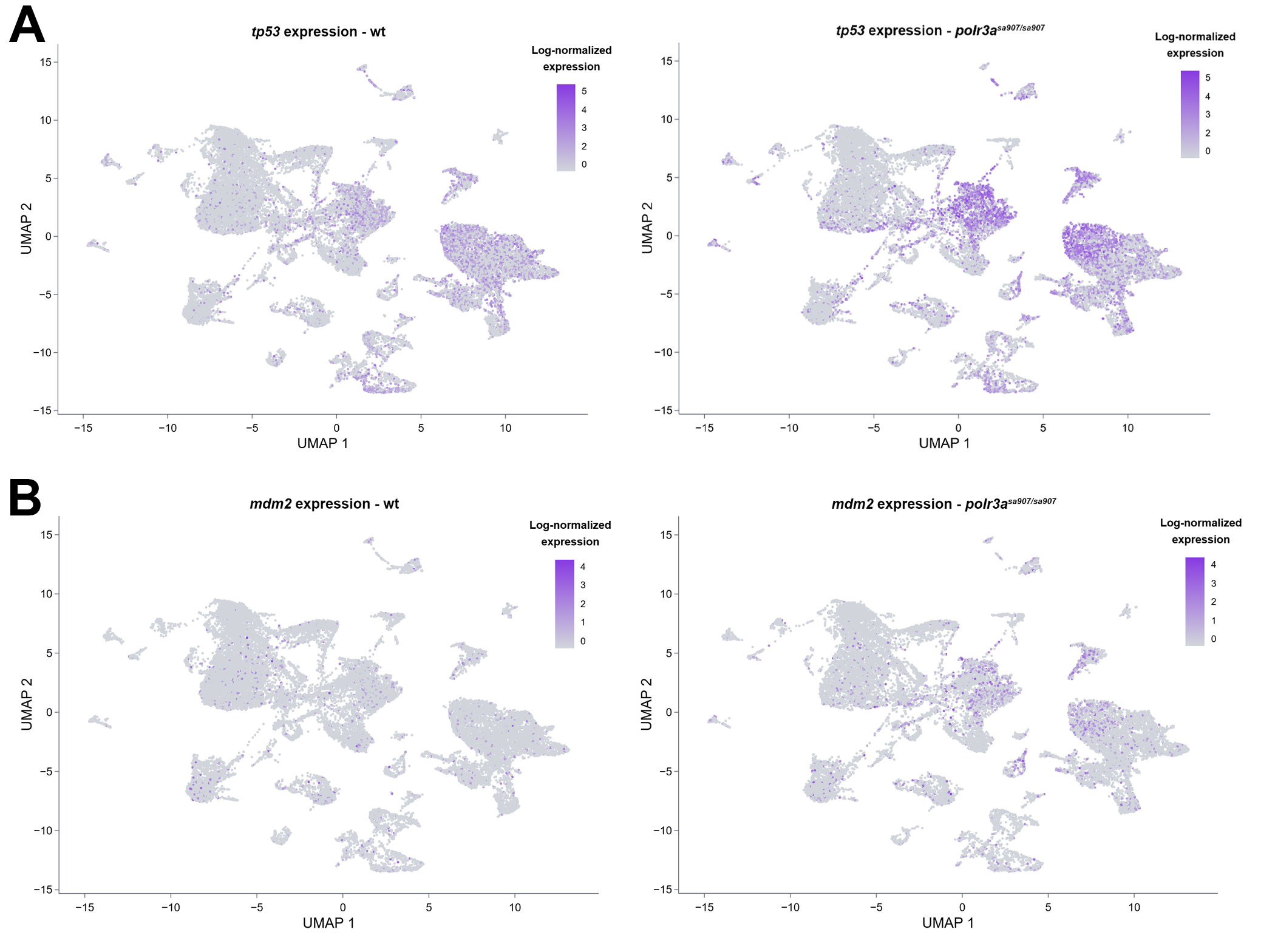

### S20 Fig

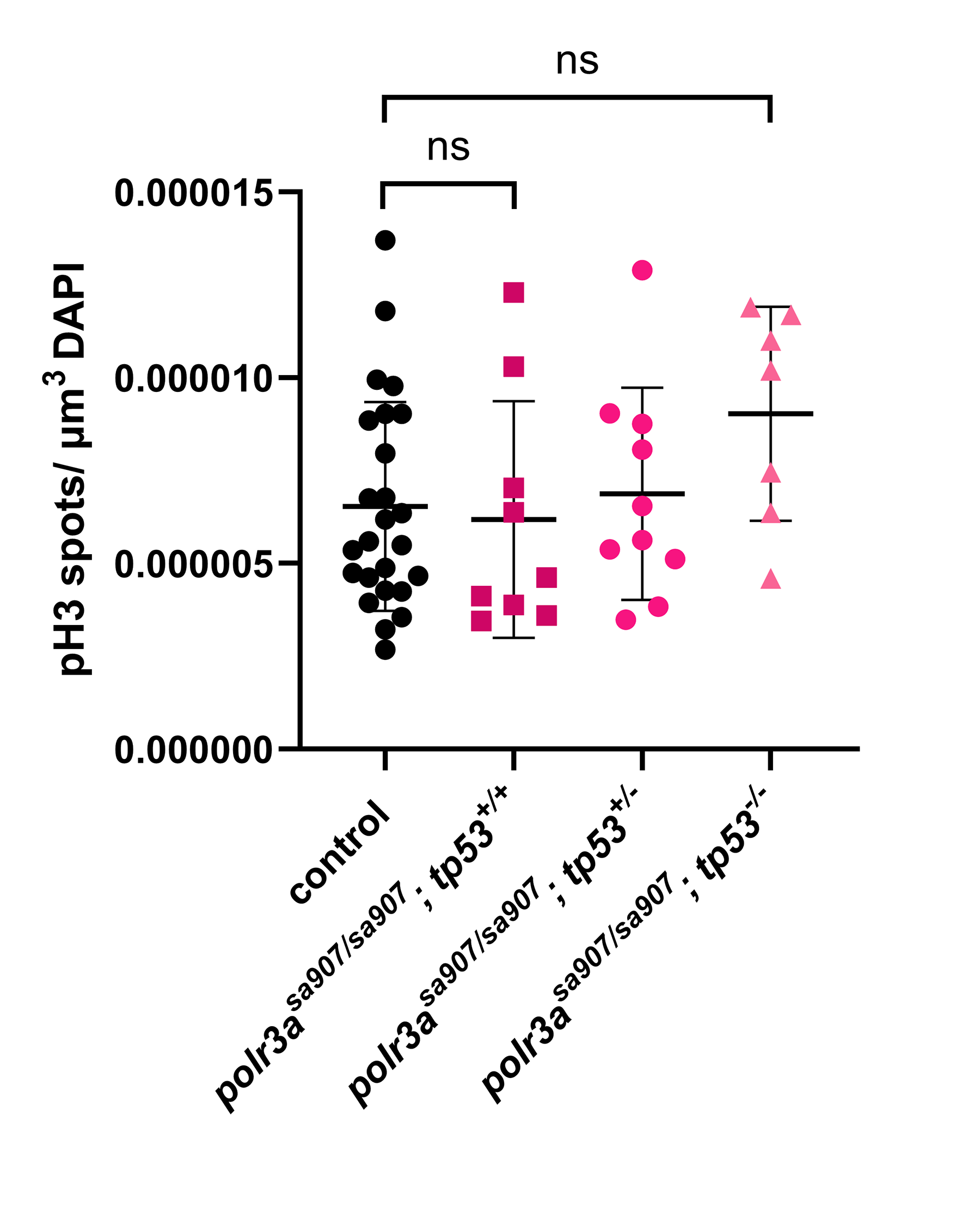

### S21 Fig

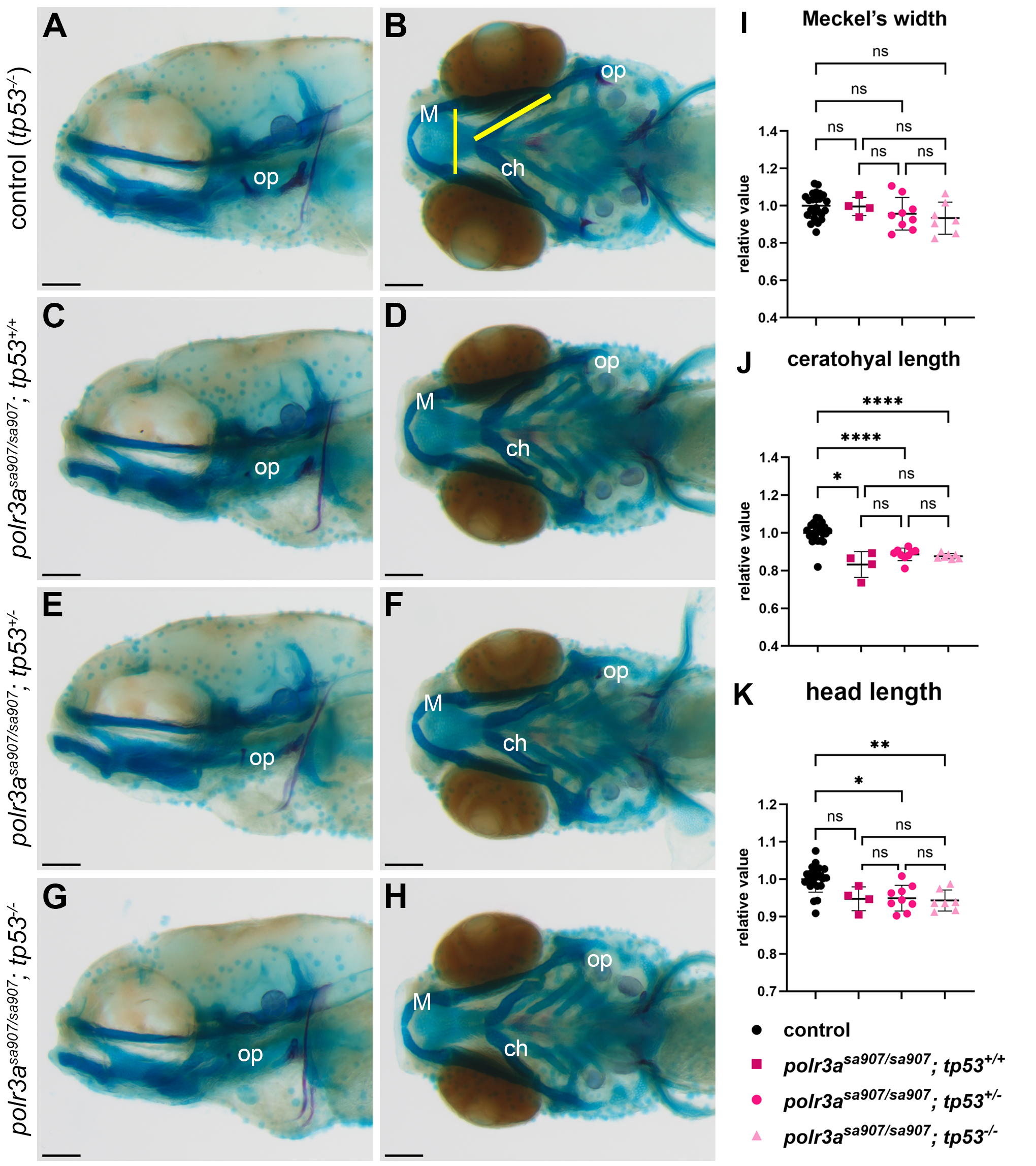

### S22 Fig

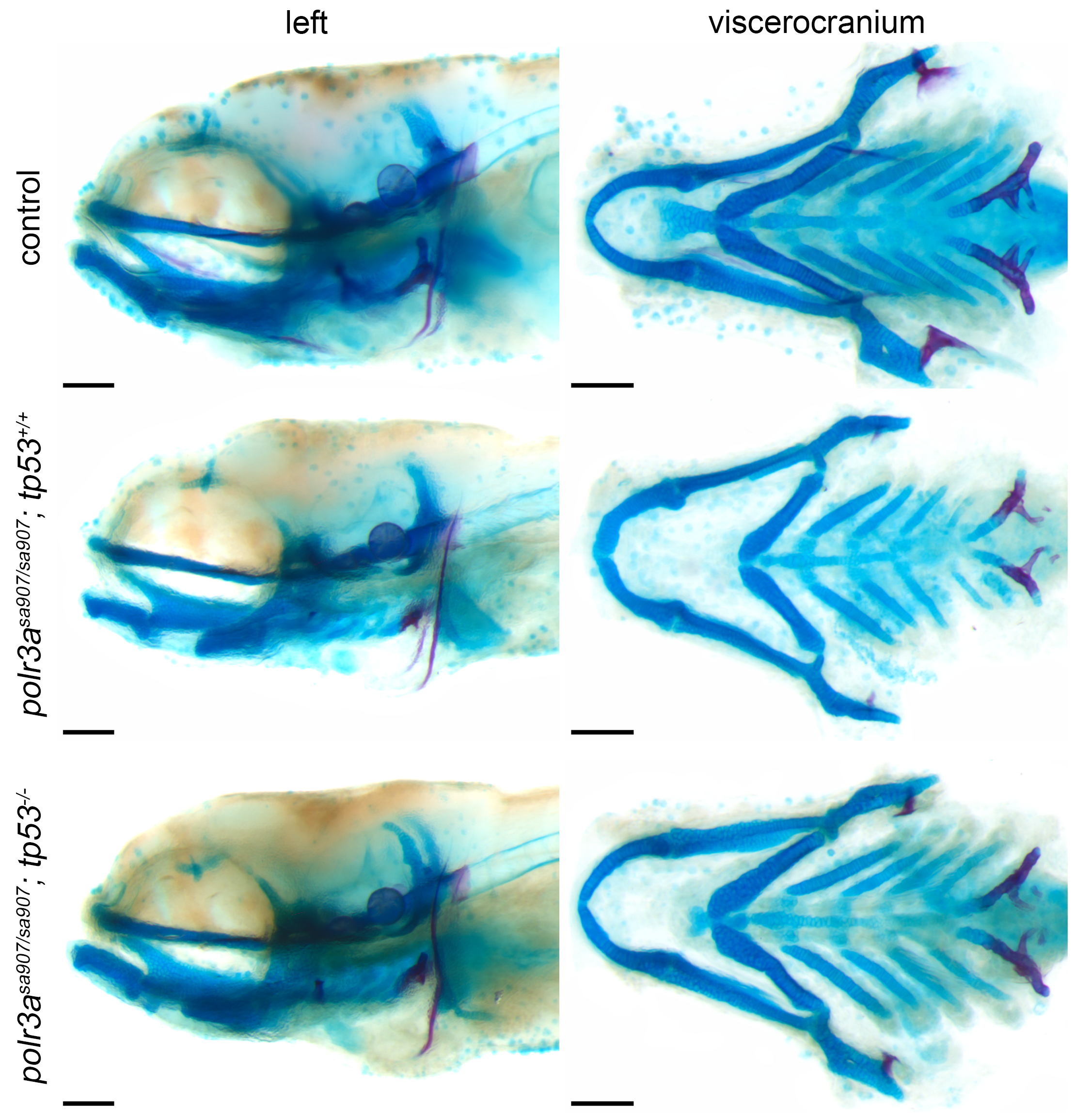
